# Genomes from 25 historical *Drosophila melanogaster* specimens illuminate adaptive and demographic changes across more than 200 years of evolution

**DOI:** 10.1101/2023.04.24.538033

**Authors:** Max Shpak, Hamid R. Ghanavi, Jeremy D. Lange, John E. Pool, Marcus C. Stensmyr

## Abstract

The ability to perform genomic sequencing on long-dead organisms is opening new frontiers in evolutionary research. These opportunities are especially profound in the case of museum collections, from which countless documented specimens may now be suitable for genomic analysis. Here, we report 25 newly sequenced genomes from museum specimens of the model organism *Drosophila melanogaster*, including the oldest extant specimens of this species. By comparing historical samples ranging from the early 1800s to 1933 against modern day genomes, we document evolution across thousands of generations, including time periods that encompass the species’ initial occupation of northern Europe and an era of rapidly increasing human activity. At the genome-wide level, we find that historical flies from the same time and place show much greater evidence for relatedness than flies from modern collections, and some show evidence of inbreeding as well, potentially reflecting either much smaller local population sizes in the past or else the specific circumstances of the collections. We also find that the Lund, Sweden population underwent local genetic differentiation during the early 1800s to 1933 interval (potentially due to accelerated drift) but then became more similar to other European populations thereafter (potentially due to increased migration). Within each time period, our temporal sampling allows us to document compelling candidates for recent natural selection. In some cases, we gain insights regarding previously implicated selection candidates, such as *ChKov1*, for which our inferred timing of selection favors the hypothesis of antiviral resistance over insecticide resistance. Other candidates are novel, such as the circadian-related gene *Ahcy*, which yields a selection signal that rivals that of the DDT resistance gene *Cyp6g1*. These insights deepen our understanding of recent evolution in a model system, and highlight the potential of future museomic studies.

## INTRODUCTION

Museum collections around the world contain billions of specimens collected over the last centuries. These collections serve as a window back in time − to an era before industrialization and modern agricultural practices − and could as such provide insights into issues ranging from insecticide resistance to climate change. The analysis of so-called historical DNA (hDNA), *i.e.* extracted genetic material from the last centuries, from museum samples accordingly holds much promise (Card et al. 2021; Raxworthy & Smith 2021). For example, whole genome analysis of historical museum samples has been used to e.g. document temporal genomic erosion in the endangered Grauer’s and mountain gorillas (van der Valk et al. 2019), and to decipher the evolutionary response of rapidly changing environmental conditions in montane chipmunks (Bi et al. 2019).

A handful of prior studies have targeted whole genome sequences from museum insect specimens. These have included proof-of-concept sequencing studies (Staats et al. 2013; Korlević et al. 2021), and investigations of the taxonomic status of museum specimens (Mikheyev et al. 2017; Grewe et al. 2021), with one study including butterfly specimens up to 150 years old (Cong et al. 2021). A few insect museomic studies have sequenced multiple individuals to examine potential targets of recent positive selection, such as focusing on the responses of honey bees to the introduction of a parasitic mite (Mikheyev et al. 2015; Parejo et al. 2020) and the emergence of the Colorado potato beetle as a crop pest (Cohen et al. 2022). In short, museomics has enabled previously unreachable research topics to be examined and brought new relevance to the myriad of specimens collected and carefully curated over the centuries in natural history collections worldwide. However, there remains much untapped potential in this area, especially in terms of sequencing larger numbers of specimens of greater antiquity, in order to extend the temporal horizon and the range of questions that museomic studies can address.

A. *D. melanogaster* has long been a primary model system for population genetics, including the identification of targets of selective sweeps. Recently, the availability of data from multiple time points has begun to improve the prospects for distinguishing natural selection from neutral evolution in this system. Genome sequencing of wild-caught and laboratory-maintained *Drosophila melanogaster* individuals has enabled the study of genomic evolution across seasonal (*e.g.* Bergland et al. 2014; Machado et al. 2021) and decadal (Lange et al. 2022) time-scales, making the species an important model for temporal genomic study. However, no previous study has investigated genome-scale variation from historical *D. melanogaster* specimens, nor from any species of such a minute animal. We here set out to sequence genomes from historical *D. melanogaster* specimens using minimally destructive techniques. Our ultimate goal was to identify changes in genome-wide diversity and specific genetic loci showing signs of having differentiated over time. Although not an agricultural pest, nor a disease vector – and accordingly not an intentional target of control methods or a casualty of altered agricultural practices – as a strict human commensal outside of its native African range, the drastic changes brought upon the planet since the dawn of the Anthropocene (here defined as the onset of the industrial revolution ca. 1780) should have left its marks in the genome of the fly. The roughly two hundred year time-frame of our analysis here should also encompass the earliest stages of this species’ adaptation to a novel high latitude environment (Keller 2007). Hence, museomic study of European *D. melanogaster* offers the potential to reveal which genes contributed to adaptive evolution during this pivotal time period. Furthermore, this interval also offers unique opportunities to investigate the demographic unknowns of early versus recent northern European *D. melanogaster* populations (such as the their local size and connectivity). Finally, for no other species do we have as detailed understanding of the genetic organization, molecular function, and the physiological and behavioral manifestations of most genes, making this fly an ideal subject for temporal genomic investigations.

The insect collections held in Lund and Stockholm are among the oldest in the world and contain many important samples, including the specimens used to erect the genus *Drosophila* (Fallén, 1814-1826). The Swedish collections span the whole Anthropocene, from the first steam engines to widespread industrialization and urbanization, to altered agricultural practices and the introduction of mass-produced chemical insecticides, to globalization and climate change (**Figure 1A**). The oldest material in the Swedish museums are four specimens from the collection of Carl Fredrik Fallén (1764-1830; **Figure 1B**) − professor in natural history at Lund university and the authority behind genus *Drosophila*. These flies are kept in the natural history museum in Stockholm but were collected in Lund. No date is provided, nor is the collector specified, but the specimens were probably collected before 1810 (see below). From about the same time stems three specimens from the collection of Johan Wilhelm Zetterstedt (1785-1874; **Figure 1C**), which are kept in Lund. Zetterstedt was a student of Fallén, and from 1822 professor in natural history at Lund University. This material comprises the lectotype (not examined here) and two paralectotypes of *Drosophila approximata*, a junior synonym of *D. melanogaster*. The two paralectotypes we have examined are labelled Lund and Smol (*i.e.* Småland) respectively, the latter indicating the province north of Scania. No collector is specified. From the collection in Copenhagen there are two specimens from Passau, Germany (**Figure 1D**) collected by Joseph Waltl (1805-1888), physician and naturalist, and from 1835 a docent and later professor in natural history at the Philosophisch-Theologische Hochschule Passau. These flies are similarly not dated but likely collected in the 1850s. Precise collection locality is unknown. Also from Copenhagen, there is one specimen from the latter half of the 19^th^ century collected by Rasmus William Traugott Schlick (1839-1916) on northeast Zealand, Denmark (**Figure 1E**). Again, no date is provided, but Schlick collected extensively in this region in the 1880s. Lastly, we have a set of 15 specimens from the collection in Lund (**Figure 1F**), collector unknown, labelled “Zootis 1933” – the informal name of the former Zoological Department at Lund University (**Figure 1G**).

**Figure 1:**
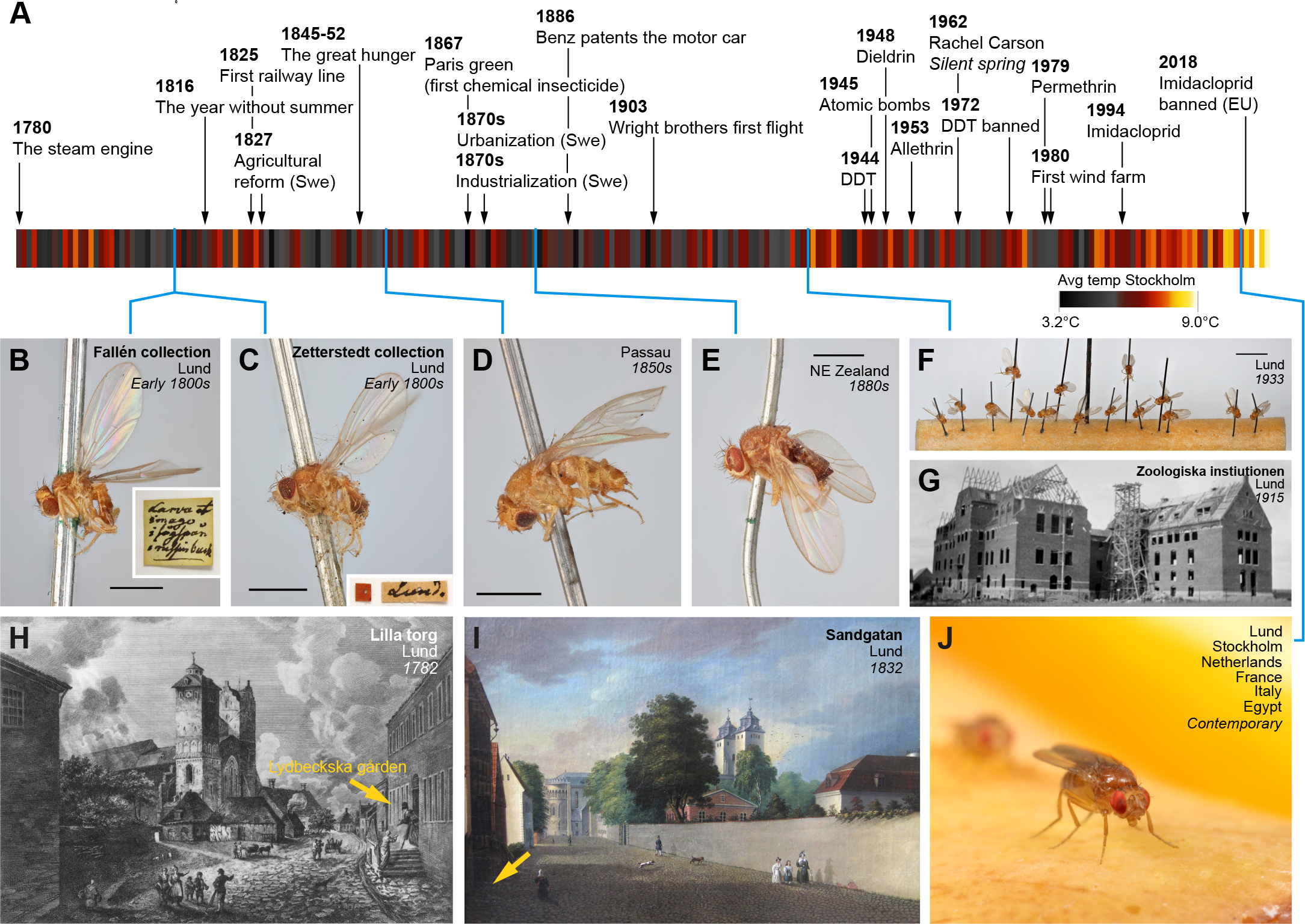
Historical context of the museum specimens and time interval encompassed by the present study. (A) Timeline of the Anthropocene showing the chronology of the analyzed fly samples, notable human events, and climate trends in Sweden (Temperature data from SMHI, Sweden). (B) One of the four *D. melanogaster* specimens from the Fallén collection at the Swedish Natural History Museum (Stockholm). Insert shows the label (in Fallén’s handwriting) accompanying the specimen. Scale bar: 1 mm. Photo: C. Fägerström. (C) One of three *D. melanogaster* specimens from the Zetterstedt collection at the Biological Museum (Lund). Insert depicts the attached label. Scale bar: 1 mm. Photo: C. Fägerström. (D) *D. melanogaster* specimen from the Natural History Museum of Denmark (Copenhagen), collected by Joseph Waltl in Passau, Germany. Scale bar: 1 mm. Photo: C. Fägerström. (E) *D. melanogaster* specimen from the Natural History Museum of Denmark (Copenhagen), collected by William Schlick on NE Själland, Denmark. Scale bar: 1 mm. Photo: C. Fägerström. (F) 15 *D. melanogaster* specimens from Lund, kept in collections of the Biological Museum (Lund). Scale bar: 5 mm. Photo: C. Fägerström. (G) The old department of Zoology building at Lund University. Photo: Lund University. (H) Lydbecksa Gården in Lund, where Fallén had his lodgnings until. Photo: Lund University. (I) Sandgatan 5 in Lund, where Zetterstedt lived. Photo: Lund University. (J) A representation of the 6 contemporary population samples included in the present analysis. Photo: M. Stensmyr

The precise timing, location, and collector for the old Swedish material is not easy to decipher. The Fallén samples are accompanied by a single (contemporary) note, in what looks like Fallén’s handwriting, stating (translated) “*Imago and larvae in sawdust [*unintelligible*] can of raisins*”. If the flies were caught by Fallén himself, a potential locality would then be his lodgings at Lidbeckska gården in central Lund (**Figure 1H**), and that the material was collected around 1807-1809, based on the known whereabouts of Fallén (who after 1811 spent considerable time at his wife’s family’s mansion at Esperöd, near Simrishamn). From his personal correspondence, the first mention of *Drosophila* can be found in a letter to Zetterstedt dated 14 September 1810 (Zetterstedt archive, Lund University). The samples in Zetterstedt’s collection are likewise difficult to precisely date. From Zetterstedt’s *Diptera Scandinaviae disposita et descripta* we can read that Zetterstedt collected *D. approximata* in Lund (possibly at his home at Sandgatan 5 in central Lund; **Figure 1I**). The specimens from Småland were attributed to Sven Ingemar Ljungh (1757-1828), civil servant and prominent naturalist, who resided at Skärsjö manor (Jönköping County, Småland – roughly 200 km from Lund). Whether or not the specimens in the collection are the ones mentioned in his treatise on Scandinavian Diptera we cannot know, but if so, the flies would likely have been caught within a decade or so of Fallén’s samples. It should be noted that Zetterstedt kept many of Fallén’s specimens in his collection, and moreover, that given the considerable age of these collections, labels might have been lost or details obscured over the years, and lastly, although both Fallén and Zetterstedt kept type specimens, detailed labelling was evidently not considered all that important.

All in all, the Swedish and Danish museum collections contain 26 *D. melanogaster* specimens suitable for analysis. The old Swedish specimens are to the best of our knowledge the earliest *D. melanogaster* available in any collection, likely predating the Meigen holotypes kept in Paris by two decades or more. We here provide whole-genome analysis from these flies, obtaining from most of them relatively complete and high quality genomes, and compare them against modern fly genomes (**Figure 1J**). This analysis may span more than 3,000 fly generations (Turelli and Hoffmann 1995; Pool 2015), analogous to studying human evolution across more than 75,000 years (approximately corresponding to the time period in which modern humans first colonized Eurasia from Africa, e.g. Mallick et al 2016, Jouganos et al 2017, Yang and Fu 2018, Wohns et al 2022). Examining changes in genomic diversity across two roughly century-scale time intervals, we find that the relationship between the Lund population and other European populations has changed over time. We identify a variety of strong candidates for the action of positive selection in each time interval, providing temporal context for previously known cases of selection, while also identifying novel putative selection targets. These findings illustrate the potential of museomic studies to deepen our understanding of recent evolution in a rapidly changing world.

## RESULTS

### Most historical flies yielded genomes of comparable quality to contemporary flies

For proof-of-principle, we first attempted to extract DNA from one of the 1933 “Zootis” specimens (**Figure 1F**). Briefly, the whole fly − still attached to the pin − was first transferred to a lysate buffer containing Proteinase K and left at 56°C for 2 hours, after which the fly was removed, washed, dehydrated, and returned to the collection (**Figure 2A**). The extraction process left the fly largely unharmed, except for a slight loss in coloration and some shrinkage of the abdomen (**Figure 2B**). The extracted DNA was subsequently prepared for Illumina short read sequencing. As expected, the DNA was highly fragmented with an average fragment length of about ∼50 bp. The Illumina run yielded ∼24 million reads, which we subsequently mapped onto a *D. melanogaster* reference genome. The ensuing historic genome was 91% complete with a 15.7X mean sequencing depth. In short, the employed method was indeed able to extract enough DNA to piece together largely complete genomes, while doing minimal harm to the specimen.

**Figure 2:**
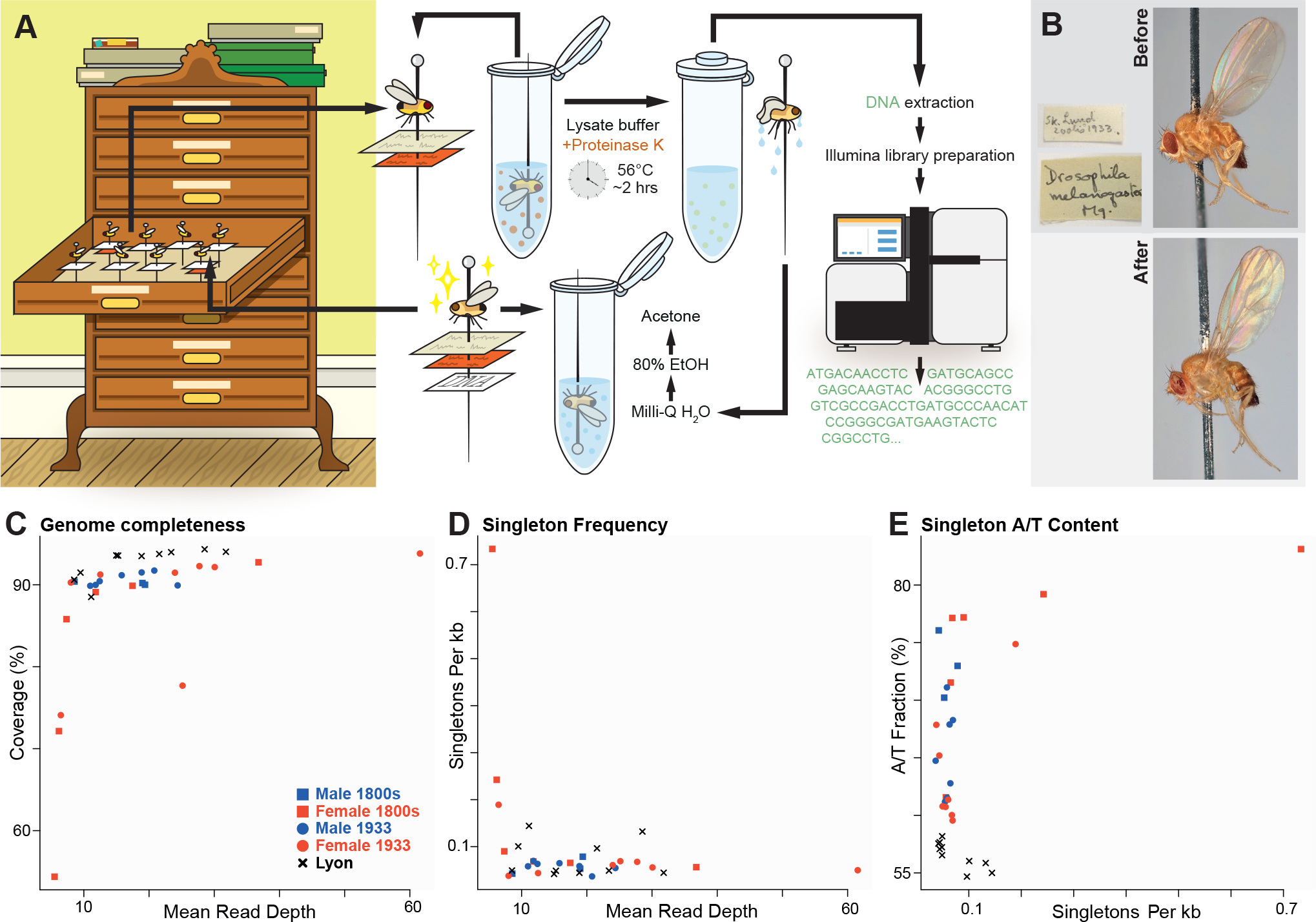
Sequencing of historical *Drosophila* specimens yields genomes with slightly reduced genomic coverage and some evidence of cytosine deamination. (A) Illustration of the process of extracting and sequencing DNA from pinned insect specimens. Flies were submerged in a lysate buffer for two hours, then put through three washing steps and dried, before being restored to the museum repository. Genomic libraries were then prepared and sequenced using the Illumina NovaSeq 6000 platform. Further details appear in the Methods section. (B) Images of the appearance of a representative specimen (from 1933) before and after the DNA extraction protocol, illustrating minor changes in coloration. Photo: C. Fägerström. (C) The fraction of genomic coverage is plotted as a function of mean per-site read depth for each historical fly genome, and for a representative set of contemporary inbred line genomes from Lyon, France. For a given level of depth most historical fly genomes achieve levels of genomic coverage only slightly lower than contemporary genomes. (D) The relationship between depth and unique (singleton) variant-calling. Only the lowest quality historical fly genomes show notable elevations in the rate of singleton variants called. (E) The fraction of singleton variants called as adenine or thymine is elevated for historical fly genomes, particularly those with greater singleton rates, potentially due to cytosine deamination.

Having verified the method, we next proceeded to process the other specimens for DNA extraction and sequencing. Out of the 25 specimens examined, we were able to generate genomes from all but one. While the mean depth of coverage per site varies among the historical fly genomes (**Figure 2C**), the median value among these genomes was approximately 20X, which is comparable to typical *D. melanogaster* genomes generated from contemporary source material (Lack et al. 2016). We find, however, that with similar mean depth, genomic coverage tends to be slightly lower in historical versus contemporary genomes (often with only 2-3% less of the genome covered; **Figure 2C**, **Table S1**).

The modest reduction in genomic coverage in most of the historical fly genomes may relate to their consistently shorter insert sizes – typically only about 50 bp (**Table S1**), compared to contemporary genomes which are typically sequenced from fragments of a few hundred bp in length. The shorter DNA fragments from the old specimens are expected to limit our ability to align into repetitive genomic regions, and hence such regions may not receive base calls in the historical genomes, leading to somewhat reduced genomic coverage as observed above. Genomes from the 1800s flies had mean insert sizes averaging 47.6 bp, while those from the 1933 collection averaged 50.8 bp. Although the distributions of mean insert sizes overlap between time points, they nevertheless differ significantly (*P* = 0.0188; Mann Whitney two-tailed *U* test). There was a marginal correlation between fragment length and depth of coverage (correlation *r* = 0.40, *P* = 0.055). A stronger and significant association was found between fragment length and genomic coverage (*r* = 0.63, *P* = 0.0001).

The shorter length of historical fly DNA fragments may be due to DNA degradation, which may also explain the reduced depth and genomic coverage obtained from a minority of the examined specimens. Among the nine 1800s samples, four genomes were characterized by a low average read depth of < 10 reads per site and/or a low genomic coverage of < 80%. For the 1933 samples, one sample was immediately excluded from further analyses due to extremely low coverage and read depth. Most samples with low coverage also had low mean read depth (**Figure 2C**; **Table S1**), and the overall correlation between these quality metrics was statistically significant (*r* = 0.404, *P* = 0.018). We note that all seven low-quality genomes (as defined above) were from female flies (out of 14 total females), whereas all 11 males yielded higher coverage/depth genomes. This sex difference is statistically significant (two-tailed binomial *P* = 0.0057) and may be due to female flies being larger and potentially experiencing greater decay before dehydration.

Regarding the six historical fly genomes retained with lower sequencing depth, inferences of diploid genotype are not statistically robust at sites for which an individual has few reads. Therefore, we treated the lower quality genomes as though they were haploid for autosomal and female X chromosomes by assigning only a single, most frequent nucleotide at each site. Consequently, there were effectively 14 rather than 18 autosome alleles for the 1800s samples and 28 rather than 30 for the 1933 autosomes. Similarly, treating each low-quality female X as haploid reduced allele counts to 11 and 21 for 1800s and 1933, respectively (from what would otherwise be X chromosome allele counts of 15 and 23).

The DNA degradation inferred above could also lead to incorrect base identification via anomalous sample-specific variants at individual sites. Therefore, we assessed whether singleton variants (unique to individual genomes) were enriched in samples with low depth and coverage. Indeed, we found a higher relative incidence of singleton variants unique to genomes with lower genomic coverage (two-tailed binomial *P* < 0.01; Figure 2B) and to a lesser extent those with low depth as well (*P* = 0.05).

One of the most common types of nucleotide degradation is cytosine deamination, which can lead to spurious thymine enrichment, which has been documented extensively in ancient DNA samples (e.g. Dabney et al 2013). We therefore assessed the extent to which A/T nucleotides are over-represented among singletons from each genome, and we found a higher fraction of A/T sites in singletons among the historical samples than in modern genomes (**Figure 2D, E; Table S1**). Whereas the modern genomes had singleton A/T frequencies of ∼55%, many of the historical genomes’ singletons were 60-74% AT, presumably due to cytosine deamination. Although most historical genomes do not show meaningfully elevated singleton rates, our findings regarding singleton AT% justify the exclusion of singleton alleles for analyses that are sensitive to rare alleles and small frequency differences, such as demographic inferences based on allele frequency change over time. In contrast, analyses that search for window-scale outliers for elevated allele frequency differentiation between samples should be little-affected by low rates of spurious singleton variants.

### The historical fly genomes show signs of relatedness and inbreeding

A preliminary analysis of pairwise genetic distances among the genomes indicated that some specific pairs of individuals from within the early 1800s and 1933 Lund samples had much greater similarity than expected. For example, as seen in **Table S2**, the raw (Hamming) distances among X chromosomes within and among modern European populations are typically 0.1-0.14. In contrast, there are 28 pairs of historical genomes with pairwise distances below 0.08 (as low as 0.038 in one case). These results suggested relatedness among some historical flies, which should lead to substantial tracts of Identity By Descent (IBD; **Figure 3A**). Because the inclusion of related individuals in estimates of allele frequency generates artifactual non-independent sampling, these redundancies should be identified and eliminated from downstream population genetic analyses.

**Figure 3:**
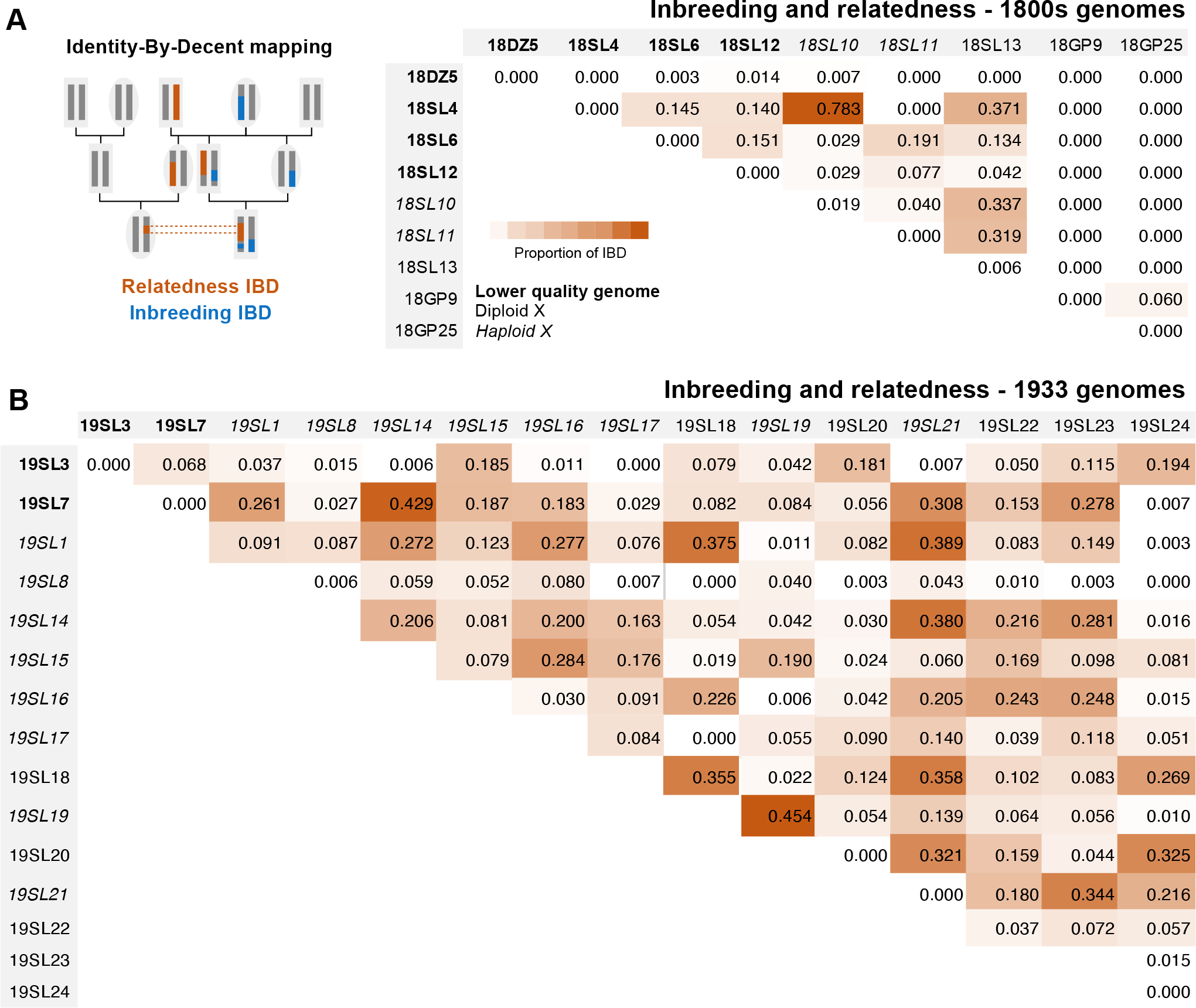
Significant levels of identity-by-descent (IBD) were observed within and between many historical fly genomes, reflecting inbreeding and relatedness. (A) Illustration of the genealogical relationships that may result in IBD due to relatedness between individuals (color1) or inbreeding within individuals (color2). (B) For the 1800s Sweden-Lund and Germany-Passau samples, the proportions of each genome detected as IBD due to apparent inbreeding (given along the diagonal) were low or zero. Above the diagonal, the proportions of each pairwise genomic comparison detected as IBD due to apparent relatedness varied within the Lund sample from zero to over 75% (the latter indicating a close familial relationship), while the two Passau genomes showed low-level relatedness IBD. (C) The same quantities are given for the Lund 1933 population sample, revealing some genomes with high levels of inbreeding, and a broad spectrum of relatedness among genomes. Since the probability of IBD detection is different for diploid and haploid sequences, males (with a single X chromsome) are given in italics. Lower quality genomes treated as haploids are shown in bold. IBD regions were masked in subsequent analyses.

We identified IBD regions by partitioning each chromosome arm into windows of approximately 100 kb length, and for each pair of individuals from the same population sample, testing whether pairwise distance in a given window was closer to simplified expectations for unrelated sequences (*i.e.* nucleotide diversity) or else IBD (*i.e.* zero for two haploid sequences, or intermediate values for IBD involving diploid sequences; see Methods). In line with the chromosome-wide distances cited above, we inferred many IBD windows spanning various lengths, up to whole chromosome arms (**Table S2; Table S3**), while **Figure 3B and C** summarize the proportion of autosomal windows with IBD between every pair of genomes in the 1800s and 1933 samples. These indicated up to sibling-level relatedness among some flies in both the 1800s and 1933 Swedish samples, and a much lower level of relatedness between the two specimens from 1840s Germany. The 1933 specimens, which are all mounted together, were likely collected at the same time, at a guess from leftover fruits on the third-floor lunch room. The six Swedish flies from the early 1800s also all show relatedness, with some pairs even at sibling-level IBD. The most plausible explanation would be that in spite of museum records tentatively linking them with three different scientists, these specimens were all from the same Lund collection event (see Discussion).

In addition to IBD between genomes, we found within-genome IBD due to inbreeding – several samples from 1933 had long regions of chromosomes with depleted heterozygosity (**Table S2; Table S4**). Two 1933 Lund samples had especially extensive regions of near-zero heterozygosity (covering 45% and 36% of their genomes; **Figure 3C**), consistent with mating between close relatives, and several other historical genomes had homozygosity across 20-40% of specific chromosome arms. Most of the genomes, however, showed no evidence of inbreeding (**Figure 3B, C**),

The pairwise and within-genome IBD indicates that many of the individuals collected for the historical were closely related and that (at least among the 1933 Lund flies) inbreeding was prevalent. These observations contrast with the low levels of relatedness observed from contemporary population samples, even when multiple fly lines founded from the same trap site were analyzed (Lack et al. 2016). The elevated IBD of the historical fly genomes could have resulted from sampling methodology and/or from a lower population density of flies in earlier eras (see Discussion).

Following IBD and inbreeding masking, we were left with 9-11 autosomal alleles for the combined 1800s samples on average rather than 14, and 10-14 rather than 28 for the 1933 samples. The average sample size of X chromosome alleles was then 8 for the 1800s set and 11 for 1933, rather than 12 and 20. Based on the mosaic pattern of window masking due to intergenomic and intragenomic IBD across individuals on each chromosome arm, post-filtering sample size varied around these averages, and downstream analyses included sample size thresholds for site inclusion (see Methods).

### Inversion frequencies appear to increase with time

We then sought to examine how within-population genetic diversity and between-population genetic differentiation have changed across more than two centuries. We first examined frequencies of known polymorphic inversions (specifically *(2L)t, (2R)Ns, (3L)P, (3R)K*, and *(3R)Mo)*, since inversions are known to significantly impact genomic diversity in this species (*e.g.* Corbett-Detig and Hartl 2012; Kapun et al. 2016). We used a previously reported list of inversion-associated SNPs (Kapun et al. 2016), a majority of which appeared to be inversion-associated in our analysis as well (see Methods and **Table S5**). Our 24 historical flies comprise a total of 192 sampled autosomal arms (as well as 37 X chromosomes), yet we inferred that only a single autosomal arm from the 1933 sample carried an inversion (being heterozygous for *(2L)t*), whereas no inversions were detected in any 1800s genome (**Table S5**). Inversion frequencies were however, also somewhat low in the modern Lund data, with only *(2L)t* (at 13%) giving a non-zero frequency estimate. The inferred frequency changes in *(2L)t* were statistically significant when modern Lund was compared to all historical genomes (Fisher exact test *P*=0.048), though not when Lund was compared to either historical sample separately (*P*=0.226 vs. 1800s and *P*=0.203 vs. 1933). These results hint at increasing inversion frequencies through time, but further temporal sampling will be needed to evaluate the previous suggestion of a recent African origin for most inversions now present in European *D. melanogaster* (Corbett-Detig and Hartl 2012; Pool et al. 2012).

### Genomic diversity suggests transiently elevated genetic differentiation

We then focused our SNP-based analysis of genetic diversity and differentiation on the X chromosome, in order to minimize the influence of inversions, which are very rare on this chromosome in European populations (*e.g.* Kapun et al. 2016). Patterns of nucleotide diversity (*π*) in Lund appeared to show a decline with time (**Figure 4A**), which could reflect the action of genetic drift. However, we note that damage-induced errors may inflate diversity estimates, especially for the 1800s sample (as potentially indicated by its slightly elevated D_xy_ values). In addition, the modern Lund sample represents a published pool-seq data set, analyzed via a distinct pipeline including the masking of rare variants, which may deflate its diversity estimate.

**Figure 4:**
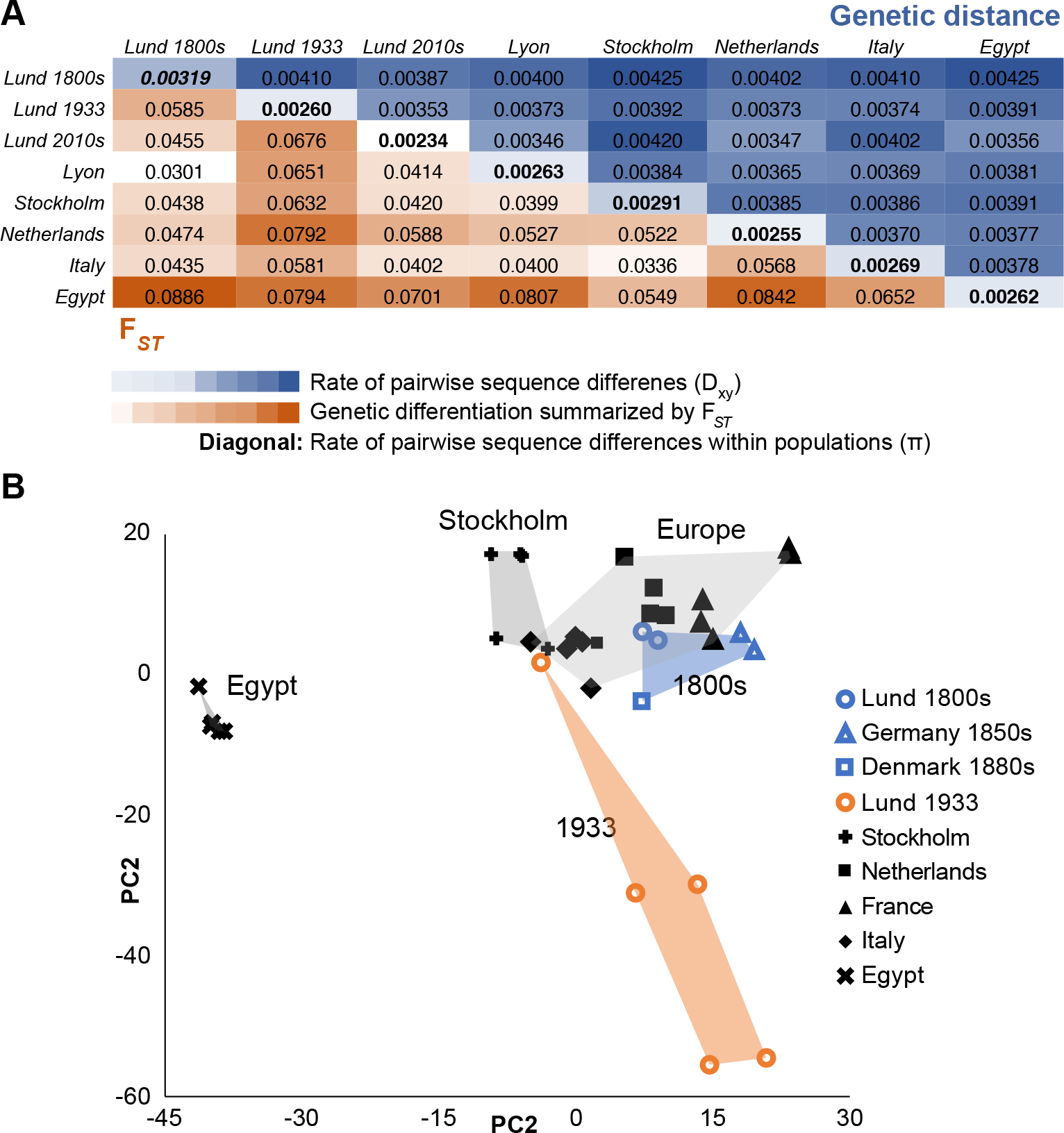
Genetic differences among historical and modern fly populations suggest transient differentiation in the Lund population. (A) Genetic differentiation between population samples is summarized by *F_ST_* (below diagonal, red heat map), and by the rate of pairwise sequence differences *D_xy_* (above diagonal, blue heat map). Along the diagonal, the rate of pairwise sequence differences within populations (nucleotide diversity) is given. Data reflects the X chromosome only (to avoid effects of inversions), with down-sampling to four alleles per population at each site. The Lund population appears to show decreasing diversity with time, subject to caveats regarding historical genome quality and the differentially-processed Lund 2010s pool-seq data. When Lund allele frequencies are compared to other European populations (via *F_ST_*), differentiation increases between the 19^th^ and 20^th^ century Lund samples, then decreases again between the 20^th^ and 21^st^ century samples. (B) Prinicple Components Analysis (PCA) results are shown for the same individually-sequenced population samples included in (A), in terms of loadings from the first two principle components. Here again, the Lund 1933 population shows elevated differentiation, whereas genomes from the three 1800s locations cluster with modern European genomes. These results reflect an analysis of X chromosome variation, with each population sample downsampled to five individuals, with non-inbred historical genomes lacking close relatedness chosen for this analysis.

Patterns of genetic differentiation (as indicated by *F_ST_*) revealed a curious temporal trajectory (**Figure 4A**). When Lund samples are compared to modern samples from around Europe, between the 1800s and 1900s sampling points, Lund becomes more differentiated from other populations. In contrast, between the 1900s and 2000s sampling points, Lund became more similar to other modern European populations. Concordant patterns were observed from Principal Components Analysis (PCA) based on X-linked variation (**Figure 4B**; **Table S6**). Here, 1800s Lund genomes were seen to cluster with modern European genomes, whereas the 1933 Lund genomes form a distinct cluster. This indication of transiently increased local genetic differentiation in Lund could indicate a history in which genetic drift led to increased allele frequency divergence from more southerly European populations, but subsequently elevated migration led to genetic homogenization between European populations (see Discussion).

We further investigated a simple model of the demographic history of Lund by using population genetic simulations to fit single-population models of genetic drift. We varied the effective population size (*N_e_*) for a given time interval and compared the mean difference in allele frequency at non-singleton SNPs between a given pair of time points from simulated versus empirical data, using a Bayesian approach to identify the best-fitting *N_e_* for each time interval, for each chromosome arm separately. In contrast to longer-term estimates of *N_e_* in this species, which are often on the order of one million (e.g. Sprengelmeyer et al. 2020), our point estimates of Lund *N_e_* were only 2,000-2,500 diploid individuals for the 1800s-1933 time interval, and 2,400-3,200 for the 1933-2010s interval (**Table S7**). These values are of the same order of magnitude as an estimate of 9,500 from a northeast United States population between 1975 and 2015 (Lange et al. 2022). Estimates from both studies share however, a key limitation in assuming that genetic drift (along with random sampling variance) is responsible for all observed frequency differences. To address how well this assumption holds, we compared *π* from coalescent simulations based on our estimated *N_e_* values to those from the empirical data, using a demographic model previously estimated for a French population (Sprengelmeyer et al. 2020) as a starting point for the pre-1800s history. While this history allowed us to match Lund ∼1809 *π* reasonably well, our low estimates of *N_e_* yielded simulated values of *π* much lower than those observed in our empirical data. For example, **Table S7** shows that for arm 3L, simulated *π* for 1933 and 2010s Lund was less than half of observed *π*. These results indicated that drift-only models did not provide an accurate approximation of the Lund population’s demographic history, and that true *N_e_* values were probably greater than our estimates. Instead, migration may have played an important role in shifting allele frequencies without decreasing *π*, as suggested above. Estimating a reasonable spatiotemporal demographic model that incorporates both local population sizes and geographic patterns of genetic structure and gene flow may require more extensive sampling of population genomic data across space and time.

### Genome scans to detect recent shifts in SNP and structural variant frequencies

One principal goal of this study is to identify instances of elevated genetic differentiation between time points that may reflect the action of recent positive selection. Our SNP-based genome scan focused on Population Branch Excess (PBE; Yassin et al. 2016), a statistic that quantifies elevated allele frequency differentiation in a focal population compared to two other reference/outgroup populations. PBE builds upon the *F_ST_*-based Population Branch Statistic (PBS; Yi et al. 2010) but is more tailored to detecting selection that is specific to the focal population. To search for selection between the early 1800s and 1933, we defined the 1800s samples as the focal population and included the Lund 1933 and Lund 2010s samples as outgroups. To focus on the 1933 to 2010s interval, Lund 1933 was the focal population and Lund 2010s and France were the outgroups. PBE was evaluated in diversity-scaled windows averaging 4.6 kb in length. In the absence of a suitable demographic model to provide a null hypothesis for the extent of genetic differentiation, we focused our analysis on the top 1% of window PBE values. However, we emphasize that any outlier-based genome scan may entail both false positives and false negatives. Our SNP-based PBE scans revealed notable outlier peaks reflecting elevated allele frequency change for each time period (**Figure 5A, B**). Overall, we identified 190 outlier regions for the 1800s-1933 time interval (referred to below as scan A) and 173 for the 1933-2010s interval (scan B), with some of these regions incorporating multiple outlier windows (**Table S8**). Since not all outliers may represent true positive targets of recent natural selection, we focus here on loci that showed the most extreme frequency changes across each century, referring to outlier regions based on their ranked maximal window PBE values.

**Figure 5:**
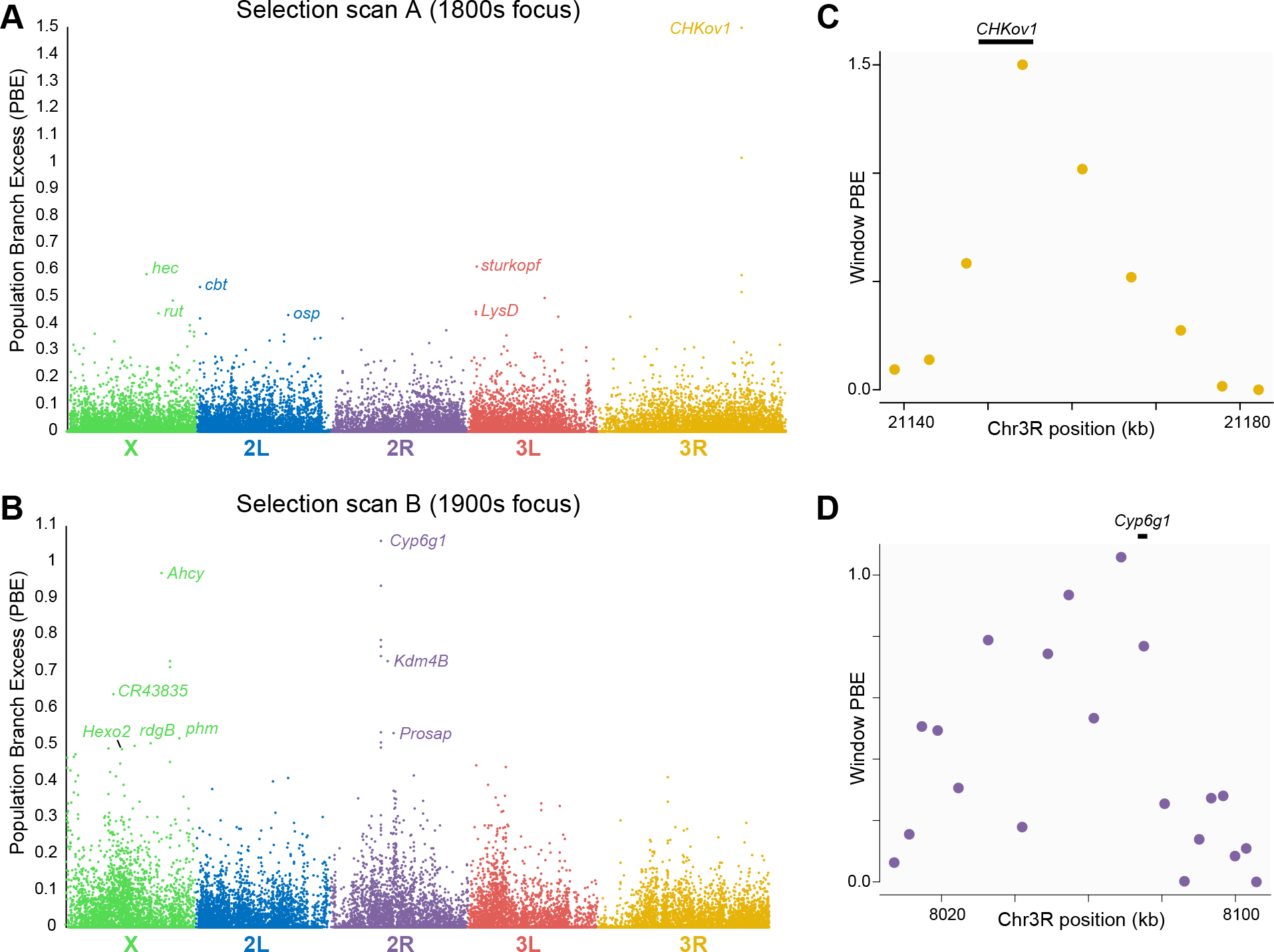
Allele frequency changes across centuries reveal candidates for positive selection. (A) Allele frequency change for ∼5 kb genomic windows, focusing on the 1800s-1933 time interval (scan A), as quantified by Population Branch Excess (PBE). Genes within top outlier regions that are discussed in the text are indicated. (B) PBE is shown for the same genomic windows, focused on the 1933-2010s time interval (scan B). (C) Window-level PBE is shown for a roughly 50 kb region focusing on the top outlier region from scan A, with the position of the known selection target *ChKov1* indicated. (D) Window-level PBE is shown for a roughly 100 kb region focusing on the top outlier region from scan B, with the position of the known selection target *Cyp6g1* indicated.

We complemented this SNP-based approach with a parallel statistical methodology to search for copy number variants (CNVs). We treated each population sample’s mean read depth within a given window (scaled relative to its chromosome arm average) as a quantitative trait, and calculated *Q_ST_* (a quantitative trait analog of *F_ST_*; Lande 1992; Spitze 1993) based on variation in read depth within and between two population samples. We then used the pairwise *Q_ST_* values among a trio of populations to calculate an analog of PBE (here referred to as QPBE), which will be elevated when the focal population has a scaled mean window sequencing depth that is much different in comparison to the two outgroup populations, potentially indicating a CNV at contrasting frequency in the focal population. It is possible that locus-specific heterogeneity in DNA degradation might also influence depth and thus QPBE values involving historical samples. Therefore, we highlight only the top result from each time interval (with further top-1% outliers given in **Table S9**), in part because both of these top outliers involve loci with previously-detected structural variants. For the 1800s to 1933 period, our top depth QPBE window contained only the gene *Tequila*, a serine protease involved in insulin/glucose regulation, lifespan, and memory (Colomb et al. 2009; Huang et al. 2015), for which polymorphic transposon insertions have created new introns (Chakraborty et al. 2018). For the 1933 to 2010s interval, the top window contained only the gene *Tenascin accessory*, involved in synapse organization (Hong et al. 2012; Mosca et al. 2012), for which polymorphic transposons are associated with expression differences (Rech et al. 2022) and balancing selection has been suggested (Croze et al. 2017). Otherwise, we refer to QPBE outliers only where they intersect with the SNP-based PBE outliers discussed below.

### Top SNP-based outliers inform classic examples of positive selection

In both time intervals, the top window PBE value was also associated with the widest outlier region, which is consistent with relatively strong positive selection. For the 1800s-1933 interval, the top outlier (labelled A1) was a 31 kb region that included the gene *CHKov1* (**Figure 5C**), which is thought to encode an ecdysteroid kinase (Scanlan *et al*. 2020). This gene is known to be associated with a protein-truncating transposon insertion under recent positive selection in non-African *D. melanogaster* populations, which was found to correlate with resistance to an organophosphate pesticide (Aminetzach *et al*. 2005). Transposon insertion and gene duplication at the *CHKov1*/*CHKov2* locus has, however, also been found to correlate with resistance to infection by the *Drosophila melanogaster* sigmavirus (DmelSV) (Magwire *et al*. 2011). Given the lack of widespread insecticide usage during this period, the timing of selection indicated by our analysis rather favor the antiviral role of this locus as a potential selective advantage. Measurements of the substitution rate in DmelSV indicate *D. melanogaster* acquired this rhabdovirus around 200 years ago (Carpenter *et al*. 2007), which predicts that this time interval should have included key stages in the evolution of resistance to DmelSV. Consistent with this hypothesis, we also found the gene *refractory to Sigma P* (*ref(2)p*) within outlier region A20. Ref2p activates the Toll pathway and has been shown to confer resistance to DmelSV infections (Bangham *et al*. 2007).

*CHKov1* and *ref(2)p* are of particular interest because mutational variants associated with viral resistance have been previously identified (Wayne et al. 1996; Magwire et al. 2011). In *CHKov1*, several SNPs have alleles in linkage disequilibrium with a transposable element insertion in the gene (Aminetzach et al. 2005) associated with viral resistance (Magwire et al. 2011). Based on seven such SNPs scored in our data, we inferred that the resistant haplotype increased from 16.7% in the 1800s sample to 100% in the 1933 and 2015 Lund samples. The occurrence of three resistance-associated haplotypes (carried in heterozygous form by two early 1800s Lund flies, as well as the later Zealand sample) could indicate that selection on this allele began prior to the collection of our 1800s flies. The fixation of the resistant haplotype in the 1933 and 2010s Lund samples suggests a complete (or nearly complete) sweep, mirroring findings from some but not all recently-collected *D. melanogaster* population samples (Aminetzach et al. 2005).

The outlier PBE score for *ref(2)p* for the 1800s time interval appeared not to be driven by the short, complex deletion identified by Wayne et al. (1996). Based on inspecting reads, this variant appeared to exist in two early 1800s Lund flies and two 1933 Lund flies, implying modest frequency change between temporal samples. The highest SNP-level PBE score at this gene was at a non-synonymous variant over 1 kb downstream from the complex deletion (2L:19544138, a threonine/serine polymorphism).

The top outlier for the 1933-2010s interval (denoted region B1) spanned 75 kb and included a known target of insecticide resistance evolution, *Cytochrome P450 6g1* (*Cyp6g1*, shown in **Figure 5D**). This gene is associated with resistance to dichlorodiphenyl-trichloroethane (DDT) and other insecticides in *D. melanogaster.* As with *ChKov1*, there is prior evidence for positive selection associated with transposon insertions and gene duplication (Daborn et al. 2002; Chung 2007; Schmidt et al. 2010; Battlay et al. 2016). Here, the novel and widespread deployment of DDT, introduced in 1944, provides a clear hypothesis for a selective pressure driving the observed frequency changes at SNPs linked to the locus in question.

### A subset of top outliers show narrow peaks of genetic differentiation

Both of the above outlier regions were relatively broad, and some of the other top outliers also showed less-broad plateaus of elevated genetic differentiation between temporal samples. In contrast, several other top outliers showed more narrowly localized signals of elevated genetic differentiation. At least in spatial analyses of local adaptation, broader intervals of genetic differentiation are more likely to be associated with selection favoring a single initial haplotype (resulting in a hard sweep), whereas narrow signals of frequency change may result from selection favoring multiple initial haplotypes, resulting in a “soft sweep” (da Silva Ribeiro et al. 2022). Narrow differentiation signals are of particular interest based on their potential to indicate not just a specific gene but often a small set of candidate variants which may include the target of selection. **Figure 6A-D** depicts four narrow SNP-level patterns of allele frequency change focused on the 1800s-1933 time interval, whereas **Figure 7A-D** depicts four such examples for the 1933-2010s interval (all of which were among the top 11 regions for window PBE for their time interval). Focusing on the earlier period, a few SNPs upstream of *hector* (*hec*; in region A3), a calcitonin-like neuropeptide receptor involved in male courtship behavior (Li et al. 2011), showed elevated PBE values (**Figure 6A**). In region A8, a cluster of SNPs within a 5’ UTR intron showed the highest PBE values at *rutabaga* (*rut*; **Figure 6B**), a calcium-responsive adenyl cyclase that influences learning, memory, lifespan, circadian rhythm, ethanol tolerance, and response to heat and oxidative stress (Levin et al. 1992; Tong et al. 2007; Donlea et al. 2009; Qin and Dubnau 2010; Scheunemann et al. 2012). We also detected a narrow set of SNPs within region A7 at the lysozyme genes *LysC* and *LysD* (**Figure 6C**), a locus for which a gene duplication and inversion polymorphisms have been reported (Chakraborty et al. 2018), and which was found to be strongly differentiated between Swedish and Italian *D. melanogaster* populations (Mateo et al. 2018). In contrast, no function is known for *CG17032* (in region A5), which also featured a cluster of high PBE SNPs (**Figure 6D**), the most extreme of which was a synonymous variant (at R5 position 3L:15953092). Among other notable 1800s-1933 outliers, the A2 region encompassed *sturkopf*, a lipid droplet-associated protein that regulates growth and stress response (Werthebach et al. 2019). A polymorphic duplication of this gene has been associated with low genetic diversity (Cardoso-Moreira et al. 2016) and this window was detected by our depth qPBE scan as well (**Table S9**). Region A4 included *cabut* (*cbt*), a transcription factor that regulates metabolic and circadian responses to nutrition (Bartok et al. 2015).

**Figure 6:**
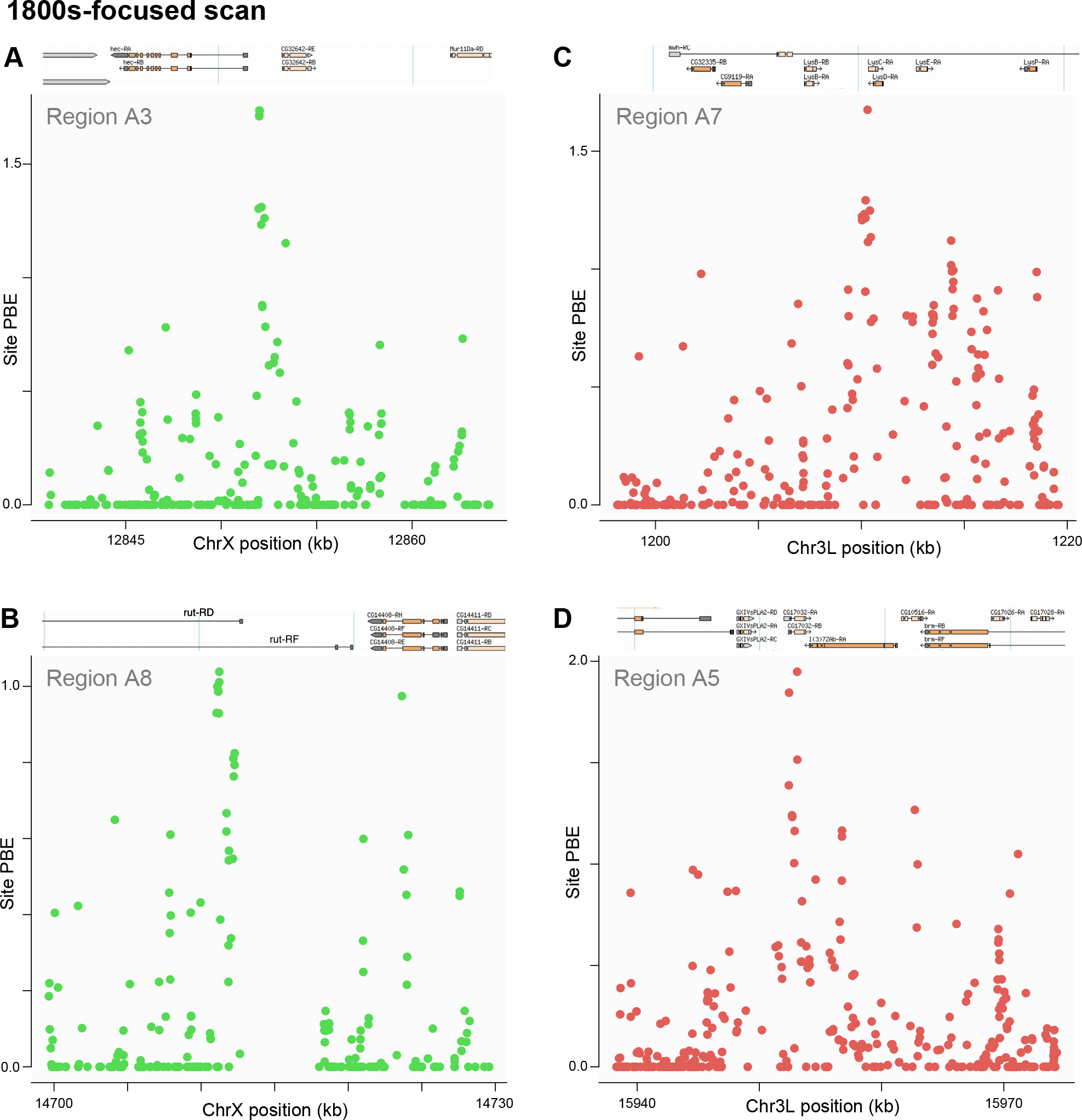
Narrow SNP-level intervals of differentiation indicate potential targets of natural selection from the 1800s-focused scan. PBE for each SNP is plotted across selected top outlier regions that showed narrow peaks of differentiation. These peaks occurred in or near: (A) *hector* (hec) in region A3, (B) *rutabaga* (rut) in region A8, (C) *Lysozyme C* (*LysC*) and *Lysozyme D* (*LysD*) in region A7, and (D) *CG17032* in region A5.

**Figure 7:**
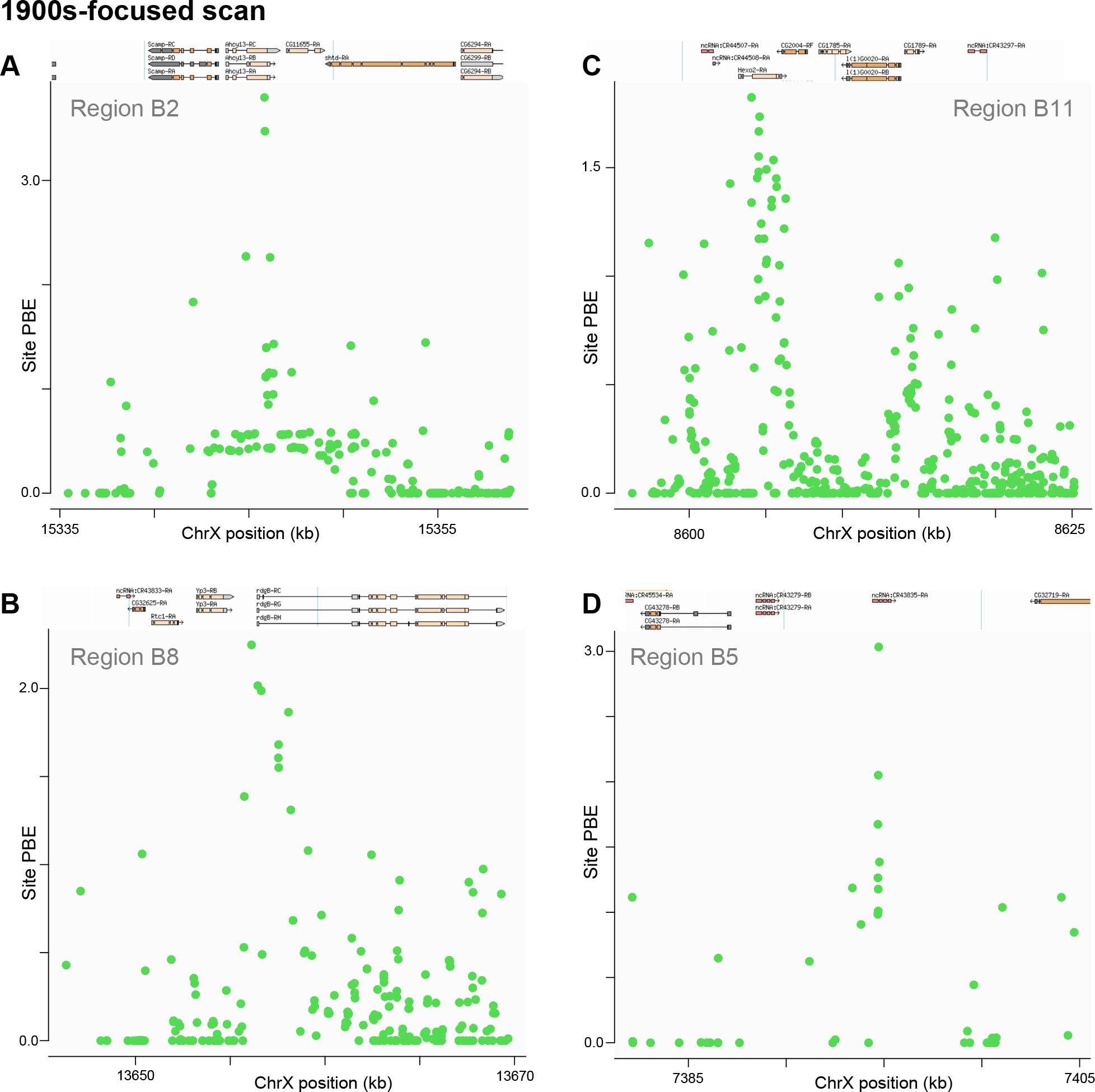
Narrow SNP-level intervals of differentiation indicate potential targets of natural selection from the 1900s-focused scan. PBE for each SNP is plotted across selected top outlier regions that showed narrow peaks of differentiation. These peaks occurred in or near: (A) *Adenosylhomocysteinase* (*Ahcy*, or *Ahcy13*) in region B2, (B) *retinal degeneration B* (*rdgB*) in region B8, (C) *Hexosaminidase 2* (*Hexo2*) in region B11, and (D) *ncRNA-CR43835* in region B5.

For the 1933-2010s interval, a window focused on *Adenosylhomocysteinase* (*Ahcy*; in region B2) yielded a PBE value nearly as extreme as that observed at *Cyp6g1*, but with a far narrower genomic scale (**Figure 7A**). Curiously, the top SNP PBE value occurs within a protein-coding exon, but at a synonymous variant. *Ahcy* regulates methionine metabolism, histone methylation, and lifespan (Parkhitko et al. 2016). It is preferentially expressed in circadian-regulating pacemaker neurons (Rivas et al. 2021), and its role in circadian regulation has been confirmed in mice (Greco et al. 2020). Despite the strength of this signal, *Ahcy* does not appear to have been detected as a top selection candidate from previously published scans of contemporary genomic variation. The SNP with the highest site PBE in *Ahcy* is associated with a synonymous variant (at R5 position 3L:15953092). We note that within the next-highest region (B3, not plotted), *Lysine demethylase 4B* (*Kdm4B*) also impacts histone methylation and circadian rhythm (Shalaby et al. 2019). For the narrow signal at region B8, the highest PBE SNPs were observed at the upstream end of *retinal degeneration B* (*rdgB*; **Figure 7B**), involved in phototransduction, olfaction, and phospholipid transport (Woodard et al. 1992; Milligan et al. 1999; Gardner et al. 2012). Notably, the narrow PBE signal at region B11 (**Figure 7C**) centered on a gene that was recently detected as a top outlier in the 1975-2015 US temporal population genomic study cited above (Lange et al. 2022); *Hexosaminidase 2* (*Hexo2*) encodes a sperm-associated protein that may play a role in fertilization (Cattaneo et al. 2006; Intra et al. 2017). Whereas another narrowly-focused top outlier (region B5) centered on the non-coding RNA gene *CR43835* (**Figure 7D**), for which no function is known.

As hinted by the exclusively X-linked examples shown in **Figure 7**, for the 1933-2010s scan specifically, there was an abundance of X-linked loci among the top outliers: although the X chromosome constitutes less than one fifth of the analyzed genome, 12 of the top 15 outlier regions were X-linked (**Table S8**). Explanations of this enrichment could include greater X chromosome effects of sampling variance (although this effect should have been stronger for the 1800s scan), genetic drift, or positive selection (potentially facilitated by X chromosome hemizygosity; Charlesworth et al. 2018). However, neither scan was meaningfully enriched for X-linked loci if the full list of outliers was considered (with the X contributing 21.7% of outliers for scan A and 18.6% for scan B). The excess of top outliers specifically for scan B could indicate that more of the X chromosome’s selection in this interval was from hard sweeps (as suggested by Harris and Garud 2023) and therefore more readily detectable from window genetic differentiation (da Silva Ribeiro et al. 2022). However, the narrower peaks of differentiation shown in **Figure 7** could indicate a role of X-linked soft sweeps as well.

### Further insights into previously implicated selection targets

Three other top outliers for the 1933-2010s interval included genes previously implicated in recent positive selection in European *D. melanogaster*. *Prosap* (in region B6) is thought to encode a structural component of the postsynaptic density (Leibl and Featherstone 2008) and affects circadian rhythm (Harbison et al. 2019). This gene was previously found to display signals of parallel adaptation across cold-adapted *D. melanogaster* populations (Pool et al. 2017), and it harbors a polymorphic transposon insertion associated with signals of selection (Rech et al. 2022). Region B7 centered on *phantom* (*phm*), which encodes a cytochrome P450 protein involved in ecdysteroid biosynthesis (Warren et al. 2004). Variation at this gene revealed a selective sweep signal in a European *D. melanogaster* population (Orengo and Aguadé 2007), and the protein displays rapid evolution between species (Good et al. 2014). Further down the list, the region B21 contained *polyhomeotic-proximal* (*ph-p*), a developmental and chromatin-modifying gene (*e.g.* Wang et al. 2006; Enderle et al. 2011), which also displayed evidence of a selective sweep in a European population (Beisswanger and Stephan 2008), associated with altered thermosensitivity of gene expression (Voigt et al. 2015). For these outliers, our findings offer insights regarding the timing of positive selection.

### Gene Ontology analysis reveals shifts in membranes and metabolism

We also applied a Gene Ontology (GO) enrichment analysis, using the permutation approach initially described in Langley et al. (2012), separately to the outliers detected from each temporal scan. From the 1800s-1933 scan, some of the most enriched categories related to cell shape and morphogenesis, including actin-mediated contraction and membrane invagination (**Table S10**). Enrichment was also observed for membrane (and vesicle) organization, compatible with the existence of population differences in membrane lipid composition (Cooper et al. 2014) and this time interval coinciding with the species’ earliest known occupation of northern Europe (Keller 2007). Curiously, “response to DDT” was also enriched, due to three cytochrome P450 outlier genes other than *Cyp6g1*, potentially reflecting pre-insecticide roles of insecticide-related genes (as with the *ChKov* locus described above). For the 1933-2010s interval, insecticide-related GO categories were not enriched among outliers. Instead, metabolic processes dominated the top results (including amide, fatty acid, lipid, and peptide metabolism), and categories related to cell division were enriched as well (**Table S10**). As with membrane composition, metabolism is also known to vary significantly between *D. melanogaster* populations from contrasting thermal environments (*e.g.* Cogni et al. 2015; Brown et al. 2018).

## DISCUSSION

Here, we report sequenced genomes from more than two dozen historical specimens of *Drosophila melanogaster*. Comparing these historical genomes to modern population genomic data, we have quantified allele frequency changes across roughly three thousand generations of evolution, and these analyses have yielded a range of insights regarding recent evolution in this model species.

### Illuminating Known and Novel Candidates for Recent Adaptive Evolution

Above, we describe genomic scans for outliers reflecting elevated SNP frequency changes across two separate time intervals (early 1800s to 1933, and 1933 to 2010s). While we report outliers reflecting the upper 1% of genomic windows for our focal statistic, we focus above on the most extreme of these outlier signals. This focus is motivated by uncertainty regarding the actual number of loci under selection in each time interval, which we do not attempt to estimate due to limitations in the data and in our ability to identify a realistic neutral model. Among the top outliers discussed above, none appeared within the full list of outliers for the alternate time period. This lack of outlier overlap could indicate that selective pressures differed between eras, or that adaptive events tended to complete within a given time period rather than spanning between them.

The strongest signals of positive selection revealed by our study added to our understanding of known selection targets, particularly with regard to the timing of their evolution. The top outlier from the 1933 to 2010s interval, *Cyp6g1*, was known to be the target of a particularly strong selective sweep related to DDT resistance, and thus provides a strong proof of concept for our ability to detect loci under recent selection. However, our inferred timing of selection for our top 1800s to 1933 outlier, *CHKov1*, served to favor one adaptive hypothesis (resistance to Sigma virus) over another (insecticide resistance). A second outlier, *ref(2)p*, reinforced the potential importance of Sigma virus as a selective pressure during our earlier time interval. As cited above, a significant proportion of our other top outliers also centered on previously reported selection targets. In addition to the above three outliers, these included *LysC*/*LysD* and *sturkopf* for the 1800s to 1933 time period, and *Hexo2*, *ph-p*, *phm*, and *Prosap* for the 1933 to 2010s interval. The number of our top outliers identified by previous studies is a testament to the depth of previous population genomic studies in *D. melanogaster*, particularly for European populations.

And yet, the majority of top outlier regions contained no previously known selection candidate. A prime example is *Ahcy* from the 1930s to 2010s scan, which showed a striking level of window frequency change comparable to *Cyp6g1*, in spite of the more narrowly focused peak at *Ahcy*. The role of *Ahcy* in circadian regulation connects it with other top outliers, especially the functionally related *Kdm4B* (the third highest outlier in this scan after *Cyp6g1* and *Ahcy*). Both *Ahcy* and *Kdm4B* alter histone methylation, an epigenetic mechanism known to regulate fly circadian rhythm (*e.g.* Shalaby et al. 2018). In addition, *Prosap* from this same 1933 to 2010s time interval, and *cbt* and *rut* from the 1800s to 1933 scan, also represent top outliers with circadian functions. The evolution of circadian behavior in high latitude *D. melanogaster* populations is well known (*e.g.* Kyriacou et al. 2008; Svetec et al. 2015). Changes in day length help regulate reproductive diapause (Saunders et al. 1989), a key adaptation in *D. melanogaster* populations from strongly seasonal climates (Schmidt et al. 2005; Erickson et al. 2020). The northerly location of Sweden entails pronounced seasonal variation in day length, which could heighten local selective pressures on daylight-responsive behaviors. While each of the above five genes have been functionally linked to circadian regulation, to our knowledge none have been linked to the evolution of circadian behavior (although *Prosap* does have documented signatures of local adaptation; Pool et al. 2017). Hence, these genes represent strong candidates for future investigations into the molecular basis of behavioral evolution.

In theory, alleles promoting adaptation to the local environment should have been adaptive in the earlier time period, and yet three of these five circadian genes were only found as outliers in the later interval. It is possible that each local population initially possessed only a subset of adaptive variants, and others arrived later via migration. Alternatively, in the 1800s to 1933 interval, the frequencies of causative variants at some circadian genes may have still been early in their rise, poising them to rise more quickly in the 1933 to 2010s interval. Another possibility is contingency – some circadian variants may have only become beneficial after other adaptive changes had occurred due to epistatic interactions among circadian genes.

### Temporal analyses reveal the complexity of demographic history

Analyses involving our museum fly genomes also yielded novel insights regarding the demographic history of a local *D. melanogaster* population from Lund, Sweden. We found that between the early 1800s and 1933, Lund genomic diversity became more differentiated from other modern European populations with respect to allele frequencies, and yet this pattern reversed between the 1933 and 2010s, with Lund regaining similarity to other European populations. This unexpected pattern of transient local differentiation suggests that distinct evolutionary forces had predominant influences on genomic diversity between these time points. As elaborated below, a primary hypothesis for this shifting pattern is that important changes in population dynamics occurred between these eras.

The increased genetic differentiation of the early 1800s to 1933 period could reflect an important influence of genetic drift, which could have occurred for multiple reasons. This interval appears to reflect the earliest days of *D. melanogaster* occupying northern Europe (Keller 2007), perhaps as shifting human activities only just permitted the species to survive in this region, and hence we might predict that initial local population sizes were low. The climate during this period was also colder than at present (**Figure 1A**), which may have also limited local population sizes. In agreement with this hypothesis, Zetterstedt commented in his treatise on Scandinavian Diptera with regard to *D. melanogaster*, “*habitat in Svecia meridionali & media rarius*”, *i.e.* in Southern and Middle Sweden rare. Furthermore, the species is conspicuously absent from Carl Linnaeus’ *Fauna Svecica* (Linnaeus 1746; 1761), nor mentioned in Johann Christian Fabricius’ *Genera insectorum* (Fabricius 1776). While these factors indicate that local population size was likely low in the early 1800s, we cannot exclude an important influence of natural selection in the observed increase in genome-wide population differentiation, as European fly populations adapted to novel environments.

The hypothesized low historical population sizes of *D. melanogaster* in Lund could also contribute to the inferred prevalence of relatedness among individuals within each historical population sample, whereas flies from modern population samples generally show much less relatedness, even when multiple individuals or lines obtained from the same fly trap are analyzed (*e.g.* Lack et al. 2016). Furthermore, a notable fraction of flies from the 1933 sample show evidence of inbreeding, indicating that their parents were related to some degree. Such observations should be more likely in a relatively small and isolated local population. However, the precise circumstances of these collections are not entirely known, and details of sampling methodology (*e.g.* if multiple flies eclosing from the same piece of fruit were collected, or if flies were raised in captivity for observation) may also contribute to the relatedness and inbreeding we infer. Such uncertainties represent an inherent limitation in studying specimens that are accompanied by finite documentation.

In contrast, the genomic homogenization between Lund and other European regions that occurred between 1933 and the 2010s could reflect increased migration through time, in conjunction with reduced genetic drift in fly populations that may have grown with time. Both of those demographic changes might be predicted based on concurrent changes in human activity during the 20^th^ century. This time period featured a five-fold increase in the human population of Lund and the surrounding region, along with a warming climate (**Figure 1A**), both of which may have been conducive to population growth in this human commensal insect (and hence reduced drift), even as new challenges such as insecticides emerged. In parallel, increased human transportation, particularly the expanded shipping of fruit and other commodities, would be expected to facilitate increased long-distance dispersal of *D. melanogaster*. While these hypothesized demographic changes could be sufficient to explain the genetic homogenization of the 1933 to 2010s time period, here again, we cannot exclude a meaningful role for natural selection (such as selective sweeps spreading between regions) in contributing to reduced differentiation.

The shifting population history potentially implied by our results raises questions about the typical consideration of demographic processes in population genetic studies. Usually, data is only available from one or more modern populations, and by necessity, fairly simple neutral models of population history are fit to the data. However, we find that when genomes from multiple temporal samples are available, we see hints of important historical changes in population size and gene flow that conventional demographic inference would be unlikely to capture. How well simple demographic models can reflect key aspects of complex demographic models, and to what degree they can recapitulate the effects of complex demographies on genomic diversity, is largely unknown. Regardless, it is clear that ancient/historical genomes have strong potential to inform our understanding of the histories of natural populations, just as they have for our own species (*e.g.* Yang and Fu 2018; Wohns et al. 2022) and for other taxa (*e.g.* van der Valk et al. 2019; Cohen et al. 2022).

### Implications of sequencing historical *D. melanogaster* specimens

This project contributes to the rapidly growing field of historical genomics and the study of temporal population genomics in natural populations. To our knowledge, our samples include the oldest pinned insects from which genome sequences have been reported (extending the temporal horizon of insect museomics from the late 1800s back to the early 1800s). Overall, 96% of our specimens yielded analyzable genomes, and 73% yielded genomes of relatively high quality. These results are encouraging for future museomic studies of small invertebrates. To our knowledge, *D. melanogaster* is the smallest animal from which historical genomes have been obtained. Surprisingly, that small size may have even been an advantage. Male *D. melanogaster* are smaller than females (roughly 1 mg vs. 1.8 mg wet weight; Hillesheim and Stearns 1991), and yet all 11 males in our study yielded high quality genomes, whereas only half of the 14 females gave comparable outcomes. It is possible that less voluminous insects such as male *D. melanogaster* experience a faster rate of desiccation relative to decay, which should improve the quality and quantity of available DNA.

The genomes of these museum specimens have added to our knowledge of not only recent evolution in *D. melanogaster*, but also the specimens themselves. Beyond confirming the sex of each fly, we inferred from relatedness that the Swedish early 1800s flies, which had been linked to three different scientists and two different localities, were actually from the same time and place. These results may reflect widespread sharing of specimens among contemporary scientists, and/or imperfect record keeping through time. Whether these flies were collected by Zetterstedt (who received his doctoral degree from Lund University in 1808) or by his mentor Fallén (who was primarily active before 1810) is difficult to say. We note that all of these Swedish early 1800s flies – including the one labelled Småland – share the same type of pin (similar to the those used by Fallén). Intriguingly, the lectotype specimen (also labelled Småland) – which we were not granted permission to sequence – is, however, mounted with a distinctly different pin. The lectotype may accordingly be one of Ljungh’s flies, as described in Zetterstedt’s *Diptera Scandinaviae*, unlike the paralectotype that evidently was caught in Lund, despite the label stating otherwise. The exact collection date of the sequenced flies is unknown, although it may have occurred before 1810 (see Introduction). Given the apparent acquisition of one of these flies by Ljungh, his death in 1828 puts a latest possible bound on the collection date. These circumstances imply that the flies examined here are the oldest surviving specimens of *D. melanogaster*, predating those described by Meigen (1830). The expanse of existing knowledge regarding the biology and genome of *D. melanogaster* has greatly enhanced the insights we have been able to draw from our museum fly genomes. In turn, our results have implications for the extensive *Drosophila* research community. We have identified various loci that represent likely targets of adaptive evolution within specific recent time intervals, and in some cases, these genes have been found to impact traits relevant to known selective pressures in the recent history of *D. melanogaster* (*e.g.* circadian regulation). Hence, our study provides a wealth of candidates for functional evolution that can be investigated in subsequent laboratory studies, through leverage of the molecular, genetic, and transgenic tools available in this model system. Although museum specimens for this (or any) species are finite, further historical genomic studies in *D. melanogaster* have the potential to greatly inform the spatial and temporal dynamics of adaptive evolution and population history, broadening the range of insights that can be obtained from genomic variation in this population genetics model species.

## METHODS

### Samples and DNA extraction

Historical specimens of *Drosophila melanogaster* were sampled from the entomological collections of Biological Museum (Lund, Sweden: MZLU), Swedish Museum of Natural History (Stockholm, Sweden: NRM), and National History Museum Denmark (Copenhagen, Denmark: NHMD) (**Table S1**). DNA was extracted from the sampled specimens using the QIAamp DNA Micro kit (Qiagen). Given the fragility of these old samples, special attention was given to preserve their integrity. The samples were placed in a microtube without removing the pin. In cases where the pin was too long to fit in the microtube, or in the case the specimen was not mounted near the end, the pin was shortened. The lysis buffer was carefully added to avoid damaging the specimens due to surface tension of the buffer prior to the addition of the Proteinase K. The microtubes were incubated at 56°C for 2 hours without shaking. The buffer containing the DNA was transferred to a new microtube, and 1 mL of MilliQ water was added to the tube containing the fly to wash away the buffer. After 30 min the water was transferred to a new microtube and stored at -20°C for future DNA extraction if needed. Then the fly was washed in successive ethanol solutions of increasing concentrations (1 mL, 50%, 70%, 80%, 99%), again paying extra attention to not harm the specimen. And finally, 500 μl of acetone was added to the specimen, which was then placed to dry in a suspended position. The rest of the DNA extraction followed the protocol, except for the final elution which was performed in two steps. In the first step, 50 μl of elution buffer was added to the columns, which were left to incubate at room temperature for 15 min prior to centrifugation. For the second step, 50 μl of elution buffer was added and the columns were incubated at 70°C for 15 min prior to centrifugation.

### Library Prep and Sequencing

The quality of the extracted DNA was visually investigated on an agarose gel (1.5%). cThe library prep followed mainly the protocol of Twort *et al*. (2021). Briefly, the fragmented DNA was blunt-end repaired with T4 Polynucleotide Kinase (New England Biolabs) and then purified with the MinElute Purification Kit (Qiagen). Next followed adaptor ligation, reaction purification, and adapter fill in. The resulting products were subsequently indexed with unique dual indexes. Indexing PCR was performed in ten independent reactions to reduce amplification bias. Each PCR reaction consisted of 12-18 cycles depending on the concentration. Indexing PCR reactions were pooled prior to purification with Sera-Mag SpeedBeads™ carboxylate-modified hydrophilic (Sigma-Aldrich). An initial bead concentration of 0.5X was used to remove long fragments that were likely contaminants of fresh DNA. Libraries were selected using a 1.8X bead concentration to size select the intended library range of ∼300 bp. The resulting libraries were quantified and quality checked using the Quanti-iT™ PicoGreen™ dsDNA assay and Bioanalyzer 2100 (Agilent technologies). The final indexed libraries were pooled together prior to sequencing on an Illumina Novaseq 6000 platform at the Swedish National Genomics Institute (NGI) in Stockholm (2x150 bp, S4 flow cell). After observing that sequenced fragments only averaged about 50 bp in length, subsequent sequencing of most specimens with 2x50 bp reads was performed by the University of Wisconsin Biotechnology Center using an Illumina NovaSeq 6000 (**Table S1**).

### Genome Alignment and Variant Calling

The procedures for alignments and variant calling used to process the museum genomes largely followed the *Drosophila* Genome Nexus (DGN) pipeline as described in Lack et al. (2015). Using this pipeline with the release 5.57 *D. melanogaster* reference genome allowed newly-assembled museum genomes to be compared against published data from modern population samples of individual genomes. Modern genomes from France and Egypt were reported in the subsequent DGN publication (Lack et al. 2016). A sample from the Netherlands was also aligned in that study, and originally sequenced by Grenier et al. (2015). Population samples from Sweden (Stockholm) and Italy were sequenced by Mateo et al. (2018) and assembled using the DGN pipeline for this study. The (distinct) processing of published pooled sequencing (pool-seq) data from Lund, Sweden (Kapun et al. 2021) is described in the next section.

Following read trimming using fastp (v0.21.0, Chen et al. 2018), we followed the DGN pipeline to map, filter, and call genotypes from the sequenced museum genomes. While the pipeline is described in detail in the previously-cited papers, we note its key feature of two rounds of mapping prior to genotyping. The initial mapping of reads to the reference genome is performed with bwa -aln v.0.5.9 (Li and Durbin 2010) and stampy (Lunter and Goodson. This step is followed by indel and variant-calling using GATK (McKenna et al 2010, Depristo et al 2011). The SNPs and indels were mapped onto the (release 5) reference genome, to create a more robust target reference for a second round of alignment of the reads with bwa -aln and stampy. It is only after this second realignment and a shift of coordinates around indels to match the reference genome that genotyping for further downstream analyses was performed.

Diploid genotyping was initially performed at all sites, except for the X chromosomes from male flies. The sex of the flies in the museum samples was not identified prior to sequencing. It was identified by calculating the ratios of average read depth in the X chromosome versus the autosomes, which is expected to be approximately 1:1 in females and 1:2 in males. All perl scripts referenced in this section and elsewhere were executed on perl v5.34.0, all python scripts were executed on python 3.8.6 (except for the stampy aligner, which was run on python 2.7.18-3 to ensure compatibility), and the R scripts were run using R v.4.2.2.

For those museum genomes with a total coverage of < 80% and average per-site read depth of < 10, we concluded that diploid genotype calls could not be confidently made. These were instead treated as “pseudo-haploid” with only a single nucleotide called per site; UnifiedGenotyper (GATK, McKenna et al 2010) was run in haploid mode for all chromosomes for these low-quality genomes. When more than a single base was identified at a (pseudo-haploid) site, we retained the most frequent allele (or selected one at random in cases of a tie). For diploid genotypes, we required at least 25% of reads to represent a minor allele, otherwise the site was classified as homozygous and assigned the more frequent allele.

### Processing and Analysis of Pool-seq Data

Published pool-seq data (Kapun et al. 2021) was obtained from modern three Lund population samples: one from summer 2014 (SRR 5647735) and two from summer 2015 (SRR8439151, 8439156). Each was derived from 40 wild-caught males. For each of these samples, alignment and post-mapping processing was performed using the same procedures as for individual genomes in the first round of the DGN pipeline, and the three pool-seq sample alignments were then merged to a single bam file. Because of the aggregate, multi-individual nature of these sequence data, frequency estimation was performed using PoolSNP (Kapun et al. 2020). We imposed an additional constraint of requiring there to be at least 5 instances of a minor allele for a site to be identified as polymorphic when filtering the VCFs, in order to minimize mislabeling read base-calling errors as SNPs. Additionally (and obviously), because individual genomes cannot be identified from pool-seq data and thus could not be used to create a revised reference genome, only a single round of alignments and genotyping was performed on the pool-seq data.

Pool-seq data introduces several statistical sampling artifacts that are not an issue with individual sequencing. Specifically, the number of reads contributed by each individual varies, so that some genomes are over-represented and others are under-represented. To deal with this, we calculate an effective allele (lineage) count (Ferretti et al. 2013), i.e. the number of *j* unique lineages at a site given *n_r_* reads and n_c_ the number of chromosomes in the sample pool (*n_c_* = 240 for autosomes, *n_c_* = 120 for X chromosomes because all of the 120 flies sampled in the pool-seq study were male). The effective count *j* follows a distribution based on Stirling numbers of the second kind *S*(*n_r_*,*j*), *i.e.* the number of ways in which *n_r_* reads can be partitioned into *j* lineages. The probability of *j* independent lineages in the sample is then:

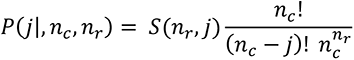

The effective number of distinct lineages (*i.e.* de facto allele count per site) was estimated from the expectation of this distribution. We found a maximum pool size of 183 for the autosomes (median = 145) and of 95 for ChrX (median = 70). Allele frequencies at each site were multiplied by the pool size to generate the appropriate down-sampled counts for population genetic analysis.

### Assessment of Genomic Data Quality

The total number of reads and the average read length was assessed from each bam file using CollectInsertSizeMetrics from the Picard suite of tools. Based on the depth and coverage criteria given above, the low-quality genomes that were treated as pseudo-haploid in the analysis pipeline were 18SL4, 18SL5, 18SL6, 18SL12 (from 1800s), 19SL3, and 19SL7 (from 1933). One sample from 1933, 19SL2, was of such low quality that it was excluded from any further processing or analysis.

To determine whether low quality sequences contributed a significant number of false positive variants in each population sample, we tallied the number of “singletons” - instances where a given genome had an allele otherwise not observed among at least five total alleles. The fraction of singleton alleles that were A or T bases was calculated to determine if cytosine deamination accounted for many of the singletons, and whether this phenomenon was correlated with sample quality. Singleton minor alleles were excluded from certain downstream analyses, such as demographic inference, while a total allele count of at least four was required for all analyses unless otherwise indicated. We also calculated the Pearson correlations between read depth and coverage (as well as between read depth and the fraction of singletons contributed by each genome, and between the fraction of singletons and the A/T content) to quantify the association between these quality metrics.

### Identification of Identity By Descent

Identity by descent (IBD) among certain genomes in both the 1800s and 1933 Lund samples was initially suspected based on highly variable pairwise differences among genomes. To identify specific regions of individual genomes with IBD, we began by partitioning chromosomes into “windows” defined by having 25,000 SNPs in an ancestral range Zambia population sample (Lack et al. 2015), averaging approximately 100 kb in length.

We then sought to account for genome-specific differences in genetic distances, such as due to sample quality. For each chromosome window of the *i*th genome, a background genetic distance *d_i_* is the median Hamming distance between the *i*th genome and unrelated modern Lyon samples. This genome-specific individual distance estimate was used to obtain a distance correction factor *m_i_* that accounts for its tendency to yield higher or lower distances than others from its population sample. The baseline population-specific distances *D* for 1800s and 1933 Lund were estimated as the median pairwise distance among individuals with no obvious IBD based on the Hamming distance matrix. Pairwise distances were compared to this baseline, scaled with respect to the Lyon outgroup distance, as follows.

For two genomes *i*,*j* collected in the same location and time point, the chromosome window Hamming distance is *d(i,j),* while *m_i_, m_j_* respectively are the scaling factors specific to sequences *i,j* – calculated from *d_i_,d_j_* divided by the median pairwise distance between all of the Lund sequences (of a given time point) and the Lyon sequences.

A. For the rescaled distance *D’(i,j) = d(i,j)m_i_m_j_*, a chromosome window is considered to be IBD between two diploid sequences (autosomes or female X chromosomes) if *D’(i,j) <* (7/8)*D*. The reasoning is that if one allele from each individual is IBD, then ¼ of pairwise allelic comparisons will yield zero distance due to this IBD, and hence the overall expected distance is ¾ of that from a non-IBD pair. For haploid to haploid (male X chromosome comparison) the same rule with a factor of ½ is applied, while for haploid to diploid (male to female X chromosome comparison) a factor of ¾ is used.

B. When pairwise IBD regions are identified, this window is masked in one genome of the pair. The masking is prioritized so that low-quality genomes are preferentially masked when compared to a higher quality genome in order to minimize data loss.

C. To avoid false negatives due to IBD regions covering partial windows, when a window was masked, the proximate half of the “upstream” and the “downstream” window was also masked. While this practice may mask some non-IBD regions, it errs on the conservative side for avoiding oversampling IBD alleles (since IBD covering more than half of a flanking window would be likely to satisfy the above masking criteria and cause the full window to be masked).

In addition to IBD among genomes, there was also evidence for depleted heterozygosity as a consequence of inbreeding in a subset of genomes from both the 1800s and (especially) the 1933 Lund populations, *i.e.* intragenomic IBD. In non-inbred genomes, the heterozygosity should approximately equal nucleotide diveristy (*π*). Therefore, to identify enrichment of homozygosity in inbred genomes, we compared the observed heterozygosity per window to the median pairwise distance among non-IBD pairs from the same population in that window. If the within-window heterozygosity of a sample was less than *π/2*, the window was flagged as IBD and a randomly selected allele of the two in the window was masked.

We included samples believed to originate from other sites and time points (18DZ5 from Zealand, Denmark and 18SL6, which was labelled as being from Smäland, Sweden) in the IBD analyses, because the latter showed low divergence when compared to some of the 1800s Lund samples. There was unambiguous evidence of close relatedness between 18SL6 and 18SL10, suggesting that the former was in fact from Lund and thus could be grouped with this population in subsequent analyses. In contrast, 18DZ5 showed roughly 1% of its genome with apparent IBD with one Lund genome, and lesser levels with two others. As these could be false positives due to conservative filtering criteria, we continued to treat this sample as being from Zealand rather than Lund in demographic analyses based on location, while at the same time masking that small region of the genome (18SL12) that showed low, IBD-like divergence with respect to 18DZ5.

### Analysis of Genetic Differentiation

Temporal and spatial divergence among different populations (defined by time and location) was assessed with analyses of population genetic distances. Our analysis focused on X chromosome sequences (to the exclusion of low recombination regions defined by Pool et al. 2017) because of the rarity of sex chromosome inversions in *Drosophila*, and only considered polymorphic sites without missing data in any of the analyzed samples.

To avoid distortions due to sample sizes, the number of alleles taken from each population (*i.e.* the 1933 Lund and the modern European and Egyptian samples) was down-sampled to match the 4 alleles from the IBD-masked 1800s Lund allele counts. This entailed selecting 4 random alleles at each site for the 1933 Lund population, the 2010s Lund pool-seq population, and the other modern populations, sampling independently at each analyzed site. This down-sampled data was used to calculate nucleotide diversity within each population, as well as between-population genetic distance (*D_xy_*) and *F_ST_* (Reynolds et al. 1983).

Genetic differentiation among the populations was also characterized using principal components analysis (PCA). Down-sampling of the Lund 1933 and modern samples was necessary to avoid over-weighting larger samples. However, because individual haplotypes were projected into the PCA space, the pool-seq Lund sample had to be excluded. To increase the number of sites with no missing data, we sampled non-masked X chromosomes that had no detectable pairwise IBD from the Hamming distance matrices from the 1800s and 1933 populations. Five such unrelated X chromosomes were identified from the 1800s samples: 18SL11, 18SL13, 18DZ5, 18GP9, and 18GP25, which were matched by the 5 unrelated 1933 Lund samples 19SL8, 19SL15, 19SL16, 19SL20, and 19SL23. Five X chromosomes were sampled from each of the modern European and Egyptian outgroup populations as well, for a total of 35 chromosomes (5 from 7 populations). For each of the 25,094 polymorphic sites with no missing data across all 35 samples in the analyzed regions of the X, the frequency of the minor allele at each site was calculated. An R-mode PCA was then used to compute the 35 eigenvectors of the correlation matrix.

### Inversion Polymorphism

To assay for inversions, we used the list of inversion-linked SNP alleles identified by Kapun (2016, 2020) for the following common autosomal inversions: *In(2L)t*, *In(2R)Ns*, *In(3L)P*, *In(3R)K*, *In(3R)Mo*, and *In(3R)P*. The frequency of each inversion was estimated from the median frequency of the inversion-associated alleles among their respective loci. For the Lund pool-seq data, the requirement of > 4 minor alleles per site to call minor alleles was relaxed to identify low-frequency inversions, *i.e.* even singleton minor alleles were included.

We used the median rather than the mean allele frequency because some alleles may be absent due to missing data. Additionally, not all of the alleles identified in Kapun (2016) as inversion-associated appeared to be fully linked to inversions in our data set. There were several instances where a small fraction of inversion-linked alleles are present in the museum samples or modern Lund (*e.g.* for *In(3R)P*, 4 of the 20 inversion-linked alleles were present at low frequency in the modern Lund pool-seq data), while the absence of inversion-linked alleles at the majority of sites suggested that the inversion itself is absent. In such cases, the use of mean allele frequency as a proxy for inversions would result in a non-zero estimated inversion frequency that is much lower than the reciprocal of the number of chromosomes sampled in the population. The estimated frequencies for each inversion in modern Lund versus the historical populations were compared using Fisher exact tests on karyotype counts, using R.

### Demographic Estimation

Using allele frequency data from 1800s, 1933, and 2010s Lund, we estimated local effective population size (*N_e_*) for the 1809-1933 and 1933-2015 intervals by implementing a simple drift-only neutral model of evolution (thus assuming no effect of migration, selection, or other processes on allele frequency changes). To estimate *N_e_* for these two time intervals, we compared the mean changes in allele frequencies across observed segregating sites to changes in allele frequencies simulated under a Fisher-Wright (FW) model of random genetic drift in discrete, non-overlapping generations. Specifically, we implemented a Bayesian approach to optimize *N_e_* that minimizes the difference between observed 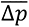, the mean absolute value difference in allele frequency across sites between 1809 and 1933 (and between 1933 and 2015) and the 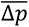 simulated under a FW model of genetic drift.

We initialized the simulations by calculating the allele frequencies at polymorphic loci at each time point in the Lund populations. Regions of low recombination, as defined by Pool et al. (2017) were excluded, as were singleton variants and non-Lund 1800s genomes. The sample frequencies *p_i_* in the 1809 population (or 1933 for the second time interval) at the *i*th locus were used to approximate the initial allele frequencies *f_i_* in the population using Bayesian estimation. The prior of *f_i_* was based on a mutation-drift equilibrium with a site mutation rate of 5.21e-10 per generation (Huang et al. 2016) and a candidate haploid effective population size *N\**. The posterior probabilities of *f_i_* were calculated prior and the observed frequencies at each site. The initial allele frequencies in each simulation replicate were generated by drawing *n* alleles equal to the number in each sample. Using a uniform prior instead resulted in very similar distributions of *f_i_*.

Population allele frequencies were initialized to *f_i_(0)* at simulation start time, representing 1809 and 1933 for the first and second time intervals, respectively. The frequencies in generation t+1 were generated by drawing a binomial sample ∼ *bin(f_i_(t), N*)* at each site, where *f_i_* = *x_i_*/*N* for *x_i_* count of what was the minor allele at *t* = 0. The model assumes a sufficiently large gene pool for sampling with replacement and treating each locus as statistically independent (*i.e.* under linkage equilibrium). This sampling is repeated for 1860 (non-overlapping) generations for 1809-1933 and 1230 generations for 1933-2015, *i.e.* for 15 generations per year (Turelli and Hoffmann 1995; Pool 2015). In the final (census) generation, an autosomal allele count *n* = 14 (*n* = 9 for ChrX) was drawn to represent the 1933 Lund sample, giving terminal sample allele frequencies *f_i_*(T)* based on *∼ bin(f_i_(T), n)*.

Much the same method was used for estimating the best fit *N_e_* for the 1933-2015 time interval, with an important exception. Because the 2010s Lund samples contributed pool-seq data, the additional sample variance that is introduced by sampling of both individuals and reads must be incorporated into the simulation. In the terminal (census) generation, the simulations implemented a two-step sampling model consistent with the nature of pool-seq data. Namely, we sampled 240 alleles (corresponding to the 120 flies) in the census generation ∼ *bin(f_i_(T), 240)*. This created a pool of 240 alleles where the frequency of the (initial) minor allele frequency was *f_i_’(T)*, with an expectation equal to *f_i_(T)*. To simulate the final sample set, we drew *n* alleles from this pool-seq subset of individuals (where *n* is the effective number of alleles per site sampled in the pool-seq data, with median values of 145 for autosomes and 70 for ChrX), *i.e. ∼ bin(f_i_’(T), n)*. This final draw gave terminal sample allele frequencies *f_i_*(T)*.

The absolute differences between final and initial sample allele frequencies in the simulations Δ*p_i_* = |*f_i_*(T)* – *p_i_(0)*| were averaged across polymorphic sites, giving 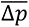. We optimized our estimate of *N_e_* by running the FW simulations over 1000 replicates over a range of *N*\*={1000…20000} in increments of 1000 and selecting the value minimizing the difference between simulated and observed 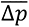. This estimate was further refined by considering *N\** in five increments of 100 on either side of the coarse-grained optimum.

We evaluated the accuracy our estimates of effective population size (*N_1_*, *N_2_*) for the time intervals 1809-1933 and 1933-2015 by comparing *π* observed in the 1800s, 1933, and 2010s Lund samples to that of simulated populations of sizes *N_1_*, *N_2_* at these time points. The simulations were initiated by sampling *N_1_* alleles from a source European population based on the demographic history of *D. melanogaster* inferred in Sprengelmeyer et al. (2020). The size *N_1_* Lund sample then experienced genetic drift (with no gene flow or natural selection) for 1860 generations (scaled in units of 4*N_e_*) until 1933, at which point the population size was set to *N_2_* and the simulations continued for a further 1230 generations. This neutral demographic model was implemented using the program ms (Hudson 1990), with the mutation rate cited above, along with a crossing-over rate of 1.09e-8, a gene conversion rate of 6.25e-8, and mean gene conversion tract length of 518 bp Comeron et al. (2012). We focused ms simulations on chromosome arm 3L because of the favorable sample sizes and relative rarity of inversion polymorphisms on this arm, and the availability of a parameterized demographic model for its evolutionary history in Europe.

### Population Branch Excess Scan for Positive Selection

To identify regions of the genome that may contain sites that may have experienced directional selection in the time between 1800s, 1900s, and modern samples, we used Population Branch Excess (PBE; Yassin et al. 2016), a statistic that aims to identify loci with particularly strong allele frequency differentiation in a focal population, in light of data from two outgroup populations. PBE is based on the Population Branch Statistic (PBS; Yi et al. 2010), which essentially estimates the length of the focal population’s branch (in terms of allele frequency differentiation). PBE is determined by a comparison of the focal population’s observed PBS value to that expected based on the lengths of the non-focal populations’ branches at the same locus, along with the average lengths of the focal and non-focal branches across loci (Yassin et al. 2016). Compared to PBS, PBE is more focused on population-specific differentiation, and will not return a large value at loci where all population branches are long (*e.g.* due to background selection or positive selection in all populations).

These statistics are based on log-transformed window *F_ST_* values *d* = -log(1-*F_ST_*) between each pair of populations. Here as in other studies, window *F_ST_* was calculated by summing the numerator terms and summing the denominator terms of the Reynolds et al. 1983 *F_ST_* estimator, and obtaining their ratio. Chromosome windows were defined to contain 250 Zambia non-singleton SNPs, corresponding to approximately 4.6 kb windows on average. We performed a separate scan for each time interval. To focus on the 1800s, we made Lund 1800s the focal population, and used Lund 1933 and Lund 2010s as the non-focal populations. To focus on the 1900s, we made Lund 1933 the focal population, and used Lund 2010s and Lyon (2010) as the outgroups. Although the Lund 2010s sample comes from (distinctly-processed) pool-seq data, the presence of one other individually-sequenced population sample among the non-focal populations should prevent any artefactual differences between individual and pool-seq data from registering as differentiation specific to the focal population.

Given that a viable demographic model was not available to provide a distribution of PBE values under a null hypothesis, outlier PBE values were defined based on the upper 1% quantile of the window distribution (which contained 240 windows genome-wide). Because strong selection can create linkage disequilibrium that extends across multiple windows, we aggregated adjacent or near-adjacent PBE-significant windows (separated by no more than 2 non-outlier windows). Although SNP-level genetic differentiation can sometimes allow the detection of soft sweeps that do not register as outlier in window-level scans, our sample sizes of museum genomes were not sufficient to enable this approach (da Silva Ribeiro et al. 2022). Instead, we used site PBE as a heuristic for identifying sites and genes of interest within window-based PBE outlier regions.

### Gene Ontology Enrichment

We performed a gene ontology (GO) enrichment analysis on genes located PBE outlier regions from the 1800s- and 1900s-focused genome scans, in order to establish whether certain functional categories (*e.g.* key metabolic or developmental pathways) were disproportionately over-represented among genes that may differ between the historical and modern populations. The GO analysis methods followed the approach of Pool et al. (2012), which involves randomly permuting the genomic locations of outlier regions, to account for variation in gene length and the genomic clustering of functionally related genes. Raw p-values were computed from 10,000 random permutations, based on the proportion of reprlicates with an equal or greater number of outlier regions associated with a given GO category as observed in the empirical data.

### Copy Number Variation

To identify potential copy number variation (CNV) with frequency differences among populations, we adopted an approach analogous to the one used above for SNP-based branch statistics. We searched for CNV differentiation among samples by comparing the mean read depth per genomic window, on the assumption that duplication (or deletion) present at elevated frequency in a given population sample will result in higher (or lower) depth of coverage, all else being equal.

Because these analyses compare read depths in museum samples with short fragments to read depths in modern samples with much longer fragments, we used trimmomatic (Bolger 2014) to reduce the read length from the modern Lyon and Stockholm samples to 50 bp. We then reran the relevant portions of the DGN assembly pipeline, while treating the truncated reads as unpaired in the mapping. We used the same genomic windows as in the SNP PBE analysis, and for each population, we calculated the mean read depth for each window across all sites in that interval. Because genomes vary in their mean depth of coverage, we corrected for such differences by dividing the mean read depth of each window by the mean depth over the entire chromosome arm. These rescaled window read depths were treated as a quantitative trait. The quantitative trait analog to the fixation index Q_ST_ (Lande 1992, Whitlock 1999), was computed for read depths among populations based on variances in read depth, as 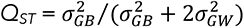, where 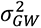 is the within-population genetic trait variance and 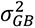 is the among-sample genetic trait variance.

By further analogy with allele frequency-based branch statistics, we calculated the quantitative trait equivalents of PBS and PBE using *d* = -log(1-*Q_ST_*) as the *Q_ST_*-based distance metric. To distinguish these trait-based branch statistics from *F_ST_*-based measures, we refer to them as QPBS and QPBE. These branch statistics were calculated for the 1800s population using 1933 Lund and modern Lyon as outgroups. For 1933 as the focal population, Lyon and Stockholm were used as the modern outgroups (since this approach is not applicable to pool-seq data such as the Lund 2010s sample). As with SNP-based PBE, we identified outlier windows in the upper 1% quantiles, and we aggregated these into regions by merging outlier windows separated by no more than one non-outlier window.

## Data Availability Statement

All new sequence data used in the present study can be found in NIH SRA Bioproject PRJNA945389. All scripts used in the present study can be found at: https://github.com/mshpak76/Museum_Drosophila_Genomes

## Acknowledgments

We wish to thank kind assistance with obtaining samples from Biologiska Museet (Lund, Sweden) and Naturhistoriska Museet (Stockholm, Sweden), and in particular Niklas Whalberg and Rune Bygebjerg. We also thank other members of our labs for providing valuable suggestions as well as relevant data and code from previous projects.

Funding was provided from Vetenskapsrådet (2019-03536) and the Max Planck Center on Next Generation Insect Chemical Ecology to MCS, and from NIH NIGMS award 144 AAH7544 to JEP.

## Author contributions

HRG and MCS collected data. MS, JDL, HRG, and JEP analyzed data. MS, JEP, and MCS wrote the manuscript.

## Supplementary Information

**Table S1. Sample Information and Genomic Sequencing Statistics.** Available collection information is given for each museum specimen. All early 1800s Sweden flies (including the one labelled Småland) were ultimately inferred to have come from the same collection event. Sequencing statistics including read count and lengths, depth of coverage, genomic coverage, singleton rate, and proportion of singletons with A or T are also given. Sample 19SL2 was excluded from most analyses because of its low sequencing depth and genomic coverage. The sex of each specimen was inferred from the relative levels of sequencing depth on the X chromosome and the autosomes. Genomic characteristics of a representative set of genomes from modern material (from Lyon, France - Lack et al. 2016 MBE) are provided, as depicted in Figure 1.

**Table S2: Evidence of identity-by-descent from relatedness and inbreeding and genomic masking proportions.**

**Tab A:** Mean pairwise genetic distance between X chromosomes from museum and modern genomes (excluding sites with missing data from any individual). Unusually low values are highlighted. Mean distance against a panel of modern genomes is indicated at the bottom.

**Tab B:** Proportion of each genome masked due to pairwise IBD with other genomes, given as a fraction of the total genome and each chromosome arm.

**Tab C:** Proportion of each genome masked due to inbreeding IBD, as indicated by runs of minimal heterozygosity, for each full genome and each chromosome arm.

**Table S3. Regions of each museum fly genome that were masked due to relatedness IBD.** Release 5 positions are given in separate tabs for each chromosome arm.

**Table S4. Regions of each museum fly genome that were masked due to inbreeding IBD.**

Release 5 positions are given in separate tabs for each chromosome arm.

**Table S5. Results of Principle Components Analysis of Genetic Structure.** Loadings for each genome are given for all 35 principle components. The X chromomsome was analyzed to avoid the influence of autosomal inversions. Up to 5 apparently unrelated individuals were included per population sample.

**Table S6. The estimated frequency of each of the most common inversions in museum and modern populations.** Inversion frequency estimates and results from each SNP reported to be associated with each inversion (by Kapun et al. 2016) are given in separate tabs.

Estimation of inversion frequency was based on the median of SNP frequencies, in order to avoid spurious non-zero estimates.

**Table S7. Effective population size estimates from drift-only models do not predict observed nucleotide diversity.**

**Top:** Mean frequency differences and haploid effective population size estimates from drift-only models for each Lund time interval.

**Bottom:** Comparison of nucleotide diversity from simulations (using estimated Ne) versus observed data, for arm 3L.

**Table S8. Outlier regions from read depth QPBE as a potential indicator of structural variants at differing frequencies between samples.**

**Tab A:** For the 1800s time interval (1800s as focal population, Lund 1933 and Lyon as outgroups), QPBE outlier regions based on the upper 1% quantile and overlapping genes are given.

**Tab B:** For the 1900s time interval (Lund 1933 as focal population, Lyon and Stockholm as outgroups), QPBE outlier regions based on the upper 1% quantile and overlapping genes are given.

**Table S9. Outlier regions from PBE reflecting windows with SNP frequencies at strongly differing frequencies between samples.**

**Tab A:** For the 1800s time interval (1800s as focal population, Lund 1933 and Lund 2010s), PBE outlier regions based on the upper 1% quantile and overlapping genes are given.

**Tab B:** For the 1900s time interval (Lund 1933 as focal population, Lund 2010s and Lyon as outgroups), PBE outlier regions based on the upper 1% quantile and overlapping genes are given.

**Table S10. Gene Ontology (GO) Enrichment based on PBE Outlier Regions. Tab A:** Gene ontology enrichment for the 1800s-focused PBE outliers.

**Tab B:** Gene ontology enrichment for the 1900s-focused PBE outliers.

For each GO category, the total number of genes in the analyzed regions is given, along with the number of outlier regions associated with these genes, and the identities of these outlier-overlapping genes. A permutation P value is also given for each GO category, which does not attempt to correct for the number of GO categories tested.

